# Conversion of cancer-associated fibroblasts from pro- to antitumor improves the sensitivity of pancreatic cancer to chemotherapeutics

**DOI:** 10.1101/2021.06.29.450327

**Authors:** Tadashi Iida, Yasuyuki Mizutani, Nobutoshi Esaki, Suzanne M. Ponik, Brian M Burkel, Liang Weng, Keiko Kuwata, Atsushi Masamune, Seiichiro Ishihara, Hisashi Haga, Kunio Kataoka, Shinji Mii, Yukihiro Shiraki, Takuya Ishikawa, Eizaburo Ohno, Hiroki Kawashima, Yoshiki Hirooka, Mitsuhiro Fujishiro, Masahide Takahashi, Atsushi Enomoto

**Author notes:** These authors contributed equally to this study (Tadashi Iida, Yasuyuki Mizutani). **Corresponding author**: Atsushi Enomoto, Department of Pathology, Nagoya University Graduate School of Medicine, 65 Tsurumai-cho, Showa-ku, Nagoya 466-8550, Japan.

## Abstract

Previous therapeutic attempts to deplete cancer-associated fibroblasts (CAFs) or inhibit their proliferation in pancreatic ductal adenocarcinoma (PDAC) were not successful in mice or patients. Thus, CAFs may be tumor suppressive or heterogeneous, with distinct cancer-restraining and -promoting CAFs (rCAFs and pCAFs, respectively). Here, we show that induced expression of the glycosylphosphatidylinositol-anchored protein Meflin, a rCAF-specific marker, in CAFs by genetic and pharmacological approaches improved the chemosensitivity of mouse PDAC. A chemical library screen identified Am80, a synthetic, non-natural retinoid, as a reagent that effectively induced Meflin expression in CAFs. Am80 administration improved the sensitivity of PDAC to chemotherapeutics, accompanied by increases in tumor vessel area and intratumoral drug delivery. Mechanistically, Meflin was involved in the suppression of tissue stiffening by interacting with lysyl oxidase to inhibit its collagen crosslinking activity. These data suggested that modulation of CAF heterogeneity may represent a strategy for PDAC treatment.

## Introduction

A well-known feature of pancreatic ductal adenocarcinoma (PDAC) is the proliferation of cancer-associated fibroblasts (CAFs), excessive deposition of the extracellular matrix (ECM) proteins produced by CAFs, and ECM remodelling in the stroma (1). CAF proliferation and ECM deposition are also found in many other recalcitrant cancers and constitute a major compartment of the tumor microenvironment (TME) along with tumor vessels and immune cells (2-7). CAFs promote cancer progression through various mechanisms and secrete various soluble factors, including growth factors, chemokines and cytokines, insoluble ECM, proteases, and extracellular vesicles, which may promote the proliferation, invasion, and metastasis of cancer cells and contribute to drug resistance and suppression of antitumor immunity (2-9). These findings have facilitated recent efforts to develop therapeutics that deplete CAFs or inhibit their proliferation and functions (2, 5-7, 10-12).

CAFs were previously thought to be a uniform population of cells with cancer-promoting functions. However, several preclinical and clinical attempts to target CAFs in PDAC mouse models and patients were not therapeutically successful, and some intervention strategies resulted in disease progression (3, 10, 11). Thus, CAFs may suppress rather than support the progression of PDAC (2, 5-7). Recent studies showed that the antitumor activity of collagen type I produced by CAFs could shape the immune microenvironment and suppress PDAC progression (13, 14). In contrast, distinct subsets of CAFs with cancer-promoting and cancer-restraining functions may exist (2, 5-7). Indeed, recent single-cell transcriptomic analyses have shown that CAFs can be classified into many subsets (15, 16). Most relevant and characterized CAF subsets in PDAC are myofibroblastic CAFs (myCAFs), inflammatory CAFs (iCAFs), and antigen-presenting CAFs (apCAFs) (16, 17). Clinicopathological and pharmacological studies have shown that these CAFs exert protumor effects through diverse mechanisms (15, 18, 19). However, the existence of cancer-restraining CAFs (rCAFs) and their marker proteins has not been investigated.

We recently identified the new CAF marker Meflin, a glycosylphosphatidylinositol (GPI)-anchored membrane protein encoded by the immunoglobulin superfamily containing leucine-rich repeat (*Islr*) gene, in both mouse and human PDAC (20, 21). Single-cell analyses of CAFs of mouse PDAC showed that Meflin marks a subset of CAFs distinct from α-smooth muscle actin (SMA)^+^ myCAFs and interleukin (IL)-6^+^ iCAFs (15, 17, 20). Furthermore, our analyses of human PDAC samples, an autochthonous PDAC mouse model, tumor transplantation models, and cell biological assays showed that Meflin suppresses PDAC progression, suggesting that Meflin is a marker of rCAFs (3, 20). Interestingly, a lineage tracing experiment showed that Meflin^+^ rCAFs, which appear in the very early stages of cancer, give rise to α-SMA^+^ CAFs that are negative or weakly-positive for Meflin during cancer progression, suggesting that Meflin^+^ rCAFs differentiate into other CAF types, including myCAFs and iCAFs (3, 20). Moreover, induction of Meflin expression in Meflin^-^ CAFs blocks PDAC progression in a tumor transplantation model, indicating that CAF function is determined by the net balance of the expression of cancer-promoting and -restraining proteins (20). In normal tissues, Meflin is expressed by pancreatic stellate cells (PSCs) in the pancreas and mesenchymal stem cells (MSCs) in the bone marrow (BM) and most other tissues, both of which are known to be major CAF origins (20-25). Therefore, rCAFs may be proliferating naïve (undifferentiated) PSCs or MSCs, which, as the tumor progresses, differentiate into Meflin-negative or weakly-positive cancer-promoting CAFs (pCAFs) (7, 20).

Given that Meflin is a functional marker of rCAFs in PDAC, one plausible strategy to treat PDAC may be to selectively induce Meflin expression in CAFs to change their phenotype. Accordingly, in this study, we first used a recombinant Sendai virus (SeV) vector to deliver the *Islr* gene, which encodes Meflin, to CAFs in a tumor transplantation mouse model. We then screened a chemical library and identified the synthetic unnatural retinoid Am80 as a compound that significantly induced Meflin expression in CAFs. Oral administration of Am80 to a PDAC transplantation mouse model significantly upregulated Meflin expression in CAFs and improved the sensitivity of tumors to conventional chemotherapeutics. We also provided mechanistic insights into the roles of Meflin in shaping the cancer-restraining TME. These findings suggest that the modulation or manipulation of CAF phenotypes by genetic or pharmacological intervention may represent a practical strategy for PDAC treatment.

## Materials and Methods

### Human tissue samples

All human PDAC samples were obtained at the time of endoscopic ultrasound-guided fine needle aspiration after the patients had provided informed consent. This study was approved by the Ethics Committee of Nagoya University Graduate School of Medicine (approval number 2017-0127-3).

### Animals

All animal protocols were reviewed and approved by the Animal Care and Use Committee of Nagoya University Graduate School of Medicine (approval numbers 30366 and 20409), and studies were conducted in compliance with institutional and national guidelines. The establishment of KPC model and Meflin-KO mice was described previously (20, 21). Genomic DNA extracted from mouse tails was used for PCR-based genotyping of WT and Meflin-KO mice. The sequences of the primers used for genotyping of the Meflin (*Islr*) gene were as follows: PCR1 forward, 5’-GCTGCATTTGAGCTGAGCCTCTGG-3’; PCR1 reverse, 5’-AACCCCTTCCTCCTACATAGTTGG-3’; PCR2 forward, 5’-TGAGGTTAGCCTGGGGACTTCAC-3’; PCR2 reverse, 5’-GGCTAGAACTCTCAAAGTAGGTCAGG-3’.

### Generation of recombinant virus, animal experiments, histology, cell biology, biochemistry, mass spectrometric analysis, and statistical analysis

Detailed protocols for the generation of recombinant virus, animal experiments, histology, cell biology, biochemistry, mass spectrometric analysis, and statistical analysis are described in **Supplementary Data**.

## Results

### Meflin expression in CAFs correlated with better responses to chemotherapeutics in human PDAC

Our previous study showed that the number of Meflin^+^ CAFs correlates with favorable clinical outcomes in patients with stage II and III borderline resectable and locally advanced PDAC (20). Here, we examined the expression of Meflin in CAFs in biopsy samples obtained from patients with PDAC who underwent chemotherapy (gemcitabine [Gem] alone, GnP [Gem plus nab-paclitaxel (nabPTX)], or FOLFIRINOX) at our institution (**Table 1**) and evaluated correlations with objective response rates (ORRs; **Fig. 1a, b**). As previously reported (20), *in situ* hybridization (ISH) analysis showed that Meflin expression was specifically observed in fibroblastic cells in the stroma, which represent CAFs, but not in other cell types such as epithelial (tumor), endothelial, smooth muscle, immune, and blood cells of PDAC tissues (**Fig. 1a**). Cases with 30% or more of stromal cells positive for Meflin were classified as Meflin high, whereas others were classified as Meflin low. Interestingly, some but not all CAFs expressed Meflin, and Meflin positivity in all CAFs was correlated with higher ORRs (**Fig. 1b**). Thus, Meflin^+^ CAFs enhanced the chemosensitivity of PDAC.

**Table 1.**
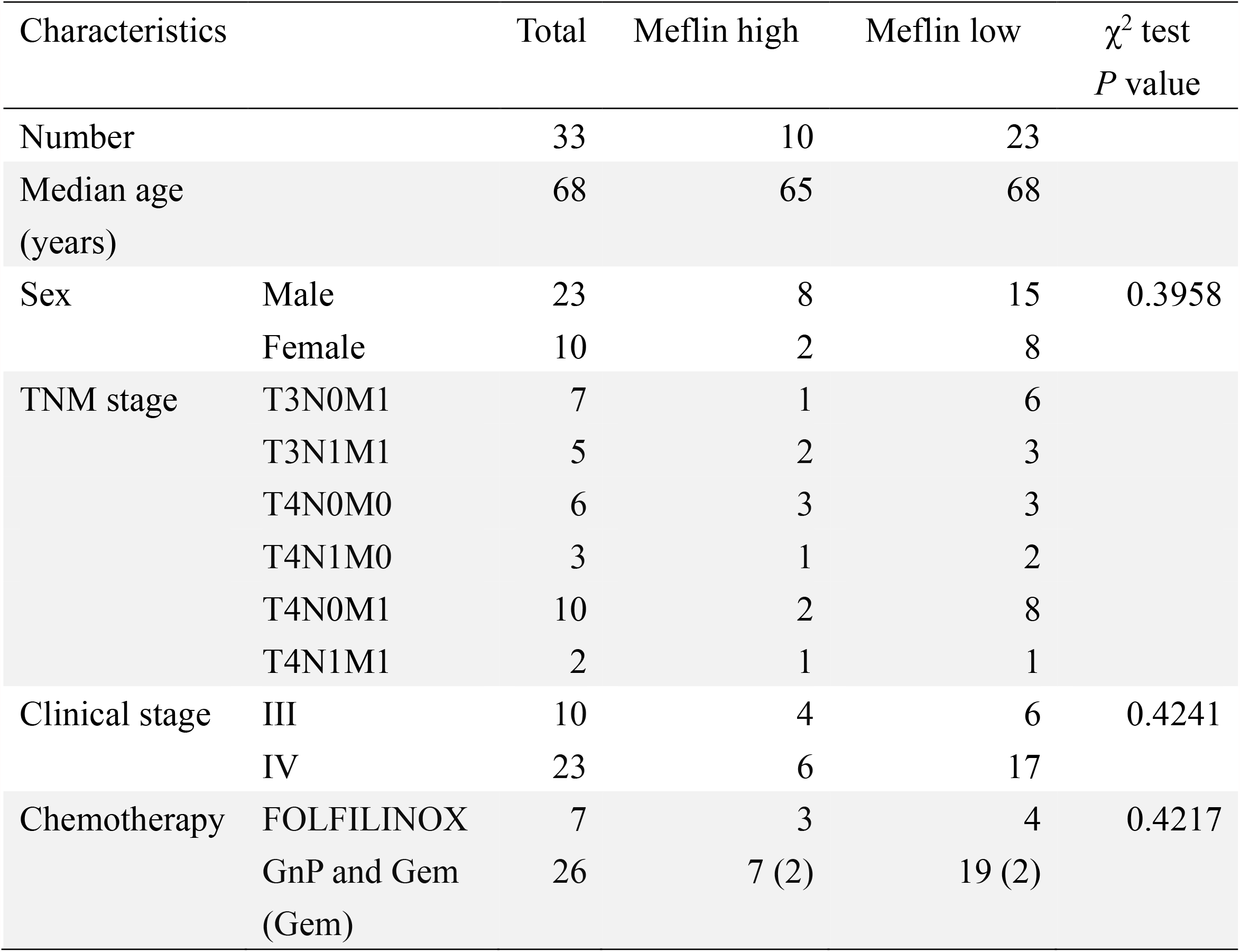
Characteristics of patients with PDAC included in the current study

**Figure 1.**
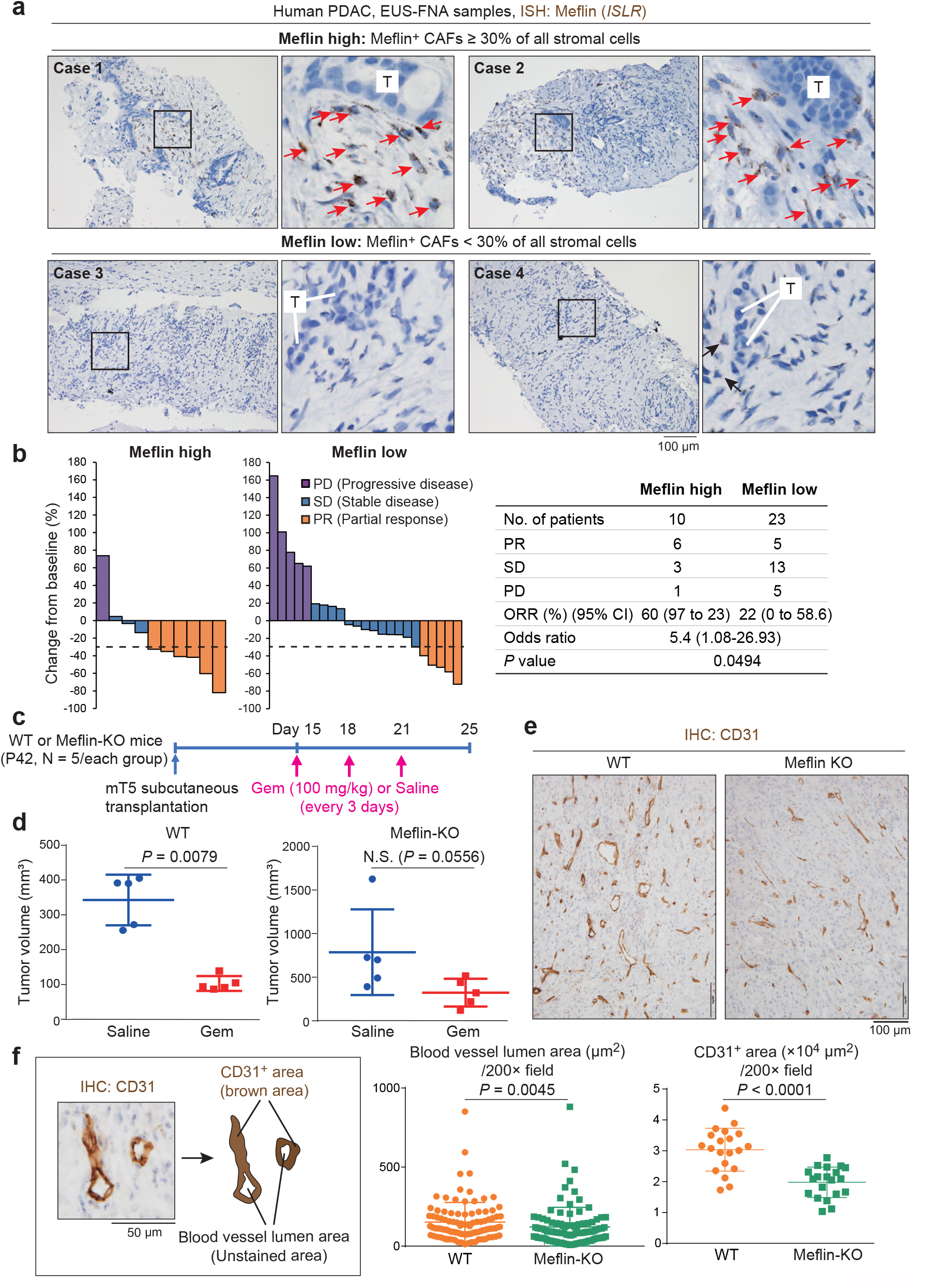
Numbers of Meflin^+^ stromal cells correlated with response of patients with PDAC to chemotherapeutics and tumor vessel area. **(a)** Representative images of Meflin ISH analysis on formalin-fixed and paraffin-embedded (FFPE) sections of biopsy samples obtained from patients with PDAC by endoscopic ultrasound-guided fine needle aspiration (EUS-FNA). Cases with 30% or more of stromal cells positive for Meflin were classified as Meflin high (cases 1 and 2, upper panel), whereas others were classified as Meflin low (cases 3 and 4, lower panel). Boxed areas were magnified in adjacent panels. Red arrows indicate Meflin^+^ CAFs. Black arrows indicate CAFs weakly positive for Meflin. T, tumor cells. **(b)** Waterfall plots depicting tumor responses of patients with PDAC (N = 33), who received Gem, nabPTX plus Gem (GnP), or FOLFIRINOX, evaluated by the Response Evaluation Criteria in Solid Tumors (RECIST) criteria. The table in the right panel shows the numbers of patients classified by their response status and Meflin expression. The results of statistical analysis are also shown. **(c)** WT and Meflin-KO female mice at postnatal day (P) 42 were subcutaneously transplanted with mT5 mouse PDAC cells (1 × 10^6^ cells/mouse), followed by i.p. administration of Gem or saline three times every 2 days and sacrifice at day 25. **(d)** Quantification of the volumes of tumors developed in WT and Meflin-KO mice administered saline (blue) or Gem (red). **(e)** Representative images of tissue sections of WT and Meflin-KO mT5 tumors stained for CD31. Brown color denotes positivity. IHC, immunohistochemistry. **(f)** Schematic illustration depicting the method of quantification of CD31-stained blood vessel area and lumen of mT5 tumors in WT and Meflin-KO mice, performed using ImageJ software (left panel). Quantification data are shown in middle and right panels.

This hypothesis was further supported by experiments in a tumor transplantation mouse model. We subcutaneously implanted mT5 cells, a mouse PDAC cell line (26), into wild-type (WT) and Meflin-knockout (KO) mice (21), followed by treatment with Gem (**Fig. 1c**). Notably, Gem treatment significantly reduced the volume of the implanted tumors in WT mice but not Meflin-KO mice, demonstrating the importance of Meflin expression in CAFs in tumor chemosensitivity (**Fig. 1d**). Consistent with our previous report showing that Meflin-KO mice developed PDAC with decreased tumor vessel areas when crossed with the Kras^LSL-G12D/+^; Trp53^LSL-R172H/+^; Pdx-1-Cre (KPC) autochthonous PDAC model (27), mT5 tumors implanted into Meflin-KO mice exhibited decreases in the lumen of tumor vessels and CD31^+^ endothelial area compared with WT mice (**Fig. 1e, f**). Thus, Meflin expression in CAFs was associated with chemosensitivity and tumor vessel perfusion, in contrast to the prevailing notion that most CAFs contribute to chemoresistance (28, 29).

### SeV-mediated Meflin expression improved PDAC sensitivity to Gem

To induce Meflin expression in CAFs, we generated a SeV encoding mouse Meflin (**Supplementary Fig. 1a, b**). Infection of primary cultured mouse MSCs with SeV-Meflin (3 × 10^7^ cell infectious units [CIU]) resulted in a significant increase in Meflin expression compared with a control SeV vector encoding DasherGFP (DGFP; **Fig. 2a**). This was accompanied by the downregulation of myCAF and iCAF markers, such as α-SMA (*Acta2*), collagen type I and III (*Col1a1 and Col3a1*), and IL6 (*Il6*; **Fig 2b**). The effects of SeV-mediated Meflin expression on CAF markers, except collagen types I and III, were also confirmed in the MSC cell line C3H10T1/2 (**Supplementary Fig. 1c**). Next, we found that injection of SeV-Meflin into subcutaneous mT5 tumors in WT mice induced Meflin overexpression without effects on α-SMA expression (**Fig. 2c, d**). Contrary to our initial expectations, we did not observe any effects of repeated transduction of SeV-Meflin on tumor progression (**Fig. 2e, f**). However, mT5 tumors transduced with SeV-Meflin exhibited a significant increase in tumor vessel area (**Fig. 2g, h**) and sensitivity to Gem treatment (**Fig. 2i, j**). The proliferation rate of mT5 cells infected with SeV-Meflin was comparable to that of control cells in culture (**Supplementary Fig. 1d**), suggesting that induction of Meflin expression in tumors enhanced their sensitivity to Gem through changes in the stromal compartment but not tumor cells.

**Figure 2.**
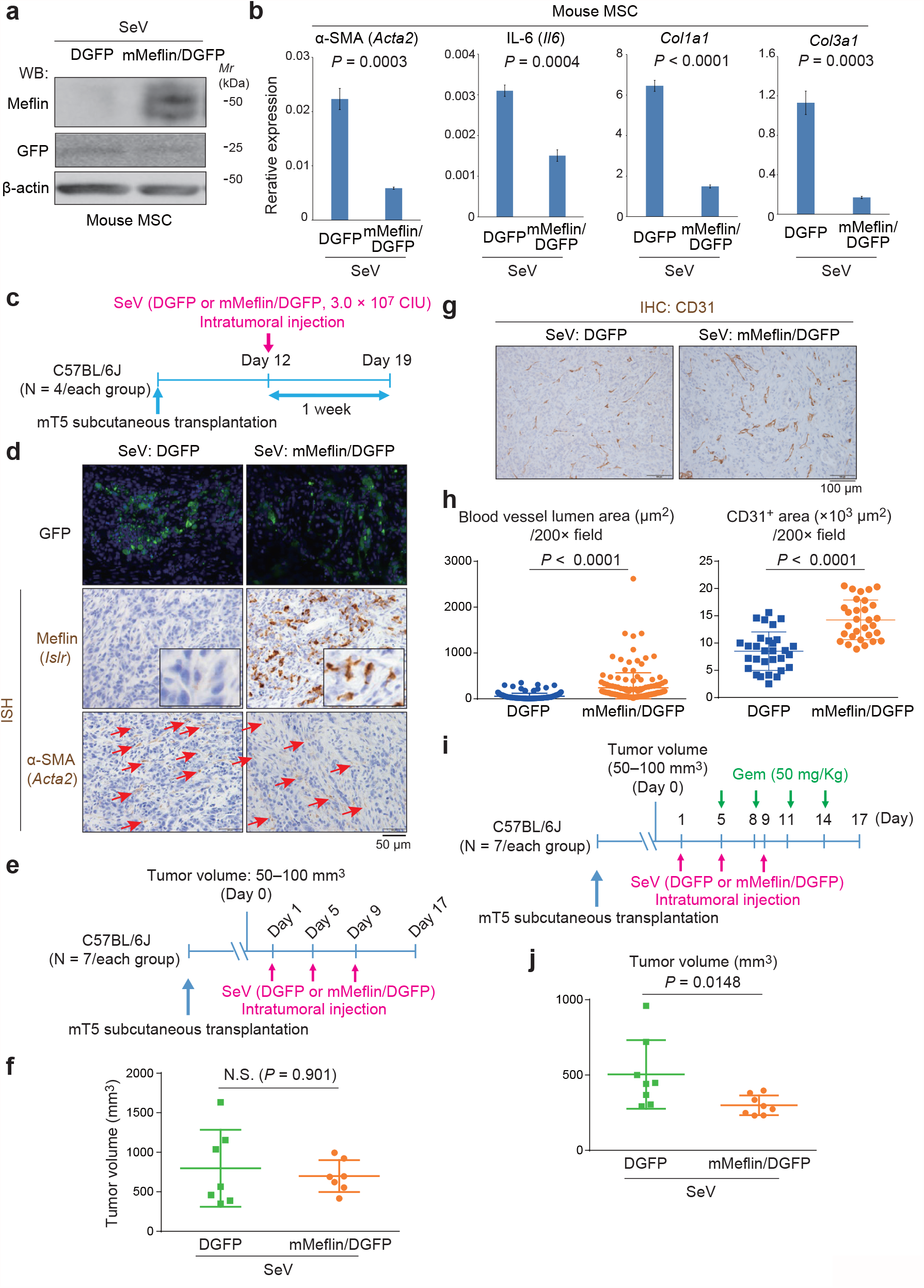
SeV-mediated transduction of Meflin induced increases in tumor vessel area and sensitivity to Gem. **(a, b)** Recombinant SeV encoding DasherGFP (DGFP) or mouse Meflin and DGFP (mMeflin/DGFP) was used to infect mouse MSCs, followed by western blot (WB) analysis with the indicated antibodies **(a)** and quantitative PCR (qPCR) for the indicated genes **(b)**. *Mr*, molecular marker. **(c)** WT female mice (P42) were subcutaneously transplanted with mT5 mouse PDAC cells (1 × 10^6^ cells/mouse), followed by intratumoral injection of the indicated SeV (3 × 10^7^ CIU) at day 12 and sacrifice at day 19. **(d)** Observation of GFP fluorescence in frozen sections prepared from mT5 tumors infected with SeV encoding DGFP (left) and Meflin/DGFP (right) (upper panel). FFPE sections were also obtained from tumors and stained for Meflin (*Islr*) and α-SMA (*Acta2*) by ISH (middle and lower panels). Arrows indicate *Acta2*^+^ CAFs. **(e–h)** mT5 tumors transplanted into WT female mice (P42) were allowed to grow until reaching 50–100 mm^3^ in volume. Mice were then injected with SeV encoding DGFP or Meflin/DGFP three times every 4 days and sacrificed on day 17 **(e)**. Tumor volume was measured **(f)**, and IHC for CD31 was performed to quantify the tumor vessel lumen and CD31^+^ area **(g, h)**. **(i, j)** mT5 tumors transplanted into WT female mice (P42) were allowed to grow until reaching 50–100 mm^3^ in volume (day 0). Mice were then injected with SeV encoding DGFP or Meflin/DGFP three times every 4 days (days 1, 5, and 9). The mice were i.p. administered Gem four times on days 5, 8, 11, and 14 and then sacrificed on day 17 **(i)**. Tumor volume was measured **(j)**.

Interestingly, we observed a rescue effect of SeV-mediated Meflin transduction on the progression of mT5 tumors developed in Meflin-KO mice (**Supplementary Fig. 2**). Induction of Meflin expression by SeV-Meflin significantly suppressed mT5 tumor growth in Meflin-KO mice, but not in WT mice (**Supplementary Fig. 2a, b**). ISH analysis showed that α-SMA expression in the tumor stroma was downregulated by Meflin expression in tumors in Meflin-KO mice (**Supplementary Fig. 2c**). These data support that Meflin may suppress α-SMA expression in CAFs and thereby block cancer progression.

### Identification of Am80 as an inducer of Meflin expression in CAFs

The findings described above suggested that programming CAFs to overexpress Meflin or convert Meflin^-/low^ CAFs to Meflin^+^ rCAFs may improve the chemosensitivity of PDAC. Moreover, administration of calcipotriol, a vitamin D analogue, to a PDAC model mice improves tumor sensitivity to Gem (30), and all-trans-retinoic acid (ATRA) reprograms activated PSCs to inhibit tumor progression through suppression of ECM remodeling (31). These studies have been the basis of ongoing clinical trials evaluating the efficacy of combination therapies of vitamin D analogues, ATRA, and conventional chemotherapeutics in patients with advanced PDAC (ref. 32; https://clinicaltrials.gov/). We previously showed that calcipotriol treatment upregulated Meflin in PDAC CAFs (20), suggesting that Meflin expression may mediate the effects of calcipotriol and ATRA on improving outcomes in patients with PDAC. Accordingly, in this study, we screened a library of nuclear receptor ligands to identify compounds that effectively upregulate Meflin expression in CAFs using human PSCs isolated from pancreatic tissues adjacent to PDAC and cultured long-term on plastic, which exhibit essentially the same properties as CAFs (30), and mouse MSCs, which are a known CAF origin in mice (**Supplementary Fig. 3a, b**). Among the compounds that upregulated Meflin in both cell types, we identified Am580, a synthetic retinoid and retinoic acid receptor (RAR) α-selective agonist (33, 34).

We next validated the effects of Am580 and another synthetic retinoid Am80, which is a retinobenzoic acid that is structurally related to Am580 (33, 34), on the expression of Meflin and other CAF marker genes in mouse MSCs (**Fig. 3a**). Am580, Am80, and ATRA upregulated Meflin and downregulated α-SMA and collagen type I and III, suggesting potential application in conversion of the CAF phenotype. Because that Am80 is approved for treating patients with recurrent and intractable acute promyelocytic leukemia (APL) in Japan (trade name: Tamibarotene) (35), we explored the *in vivo* effects of Am80 in PDAC progression in the further experiments.

**Figure 3.**
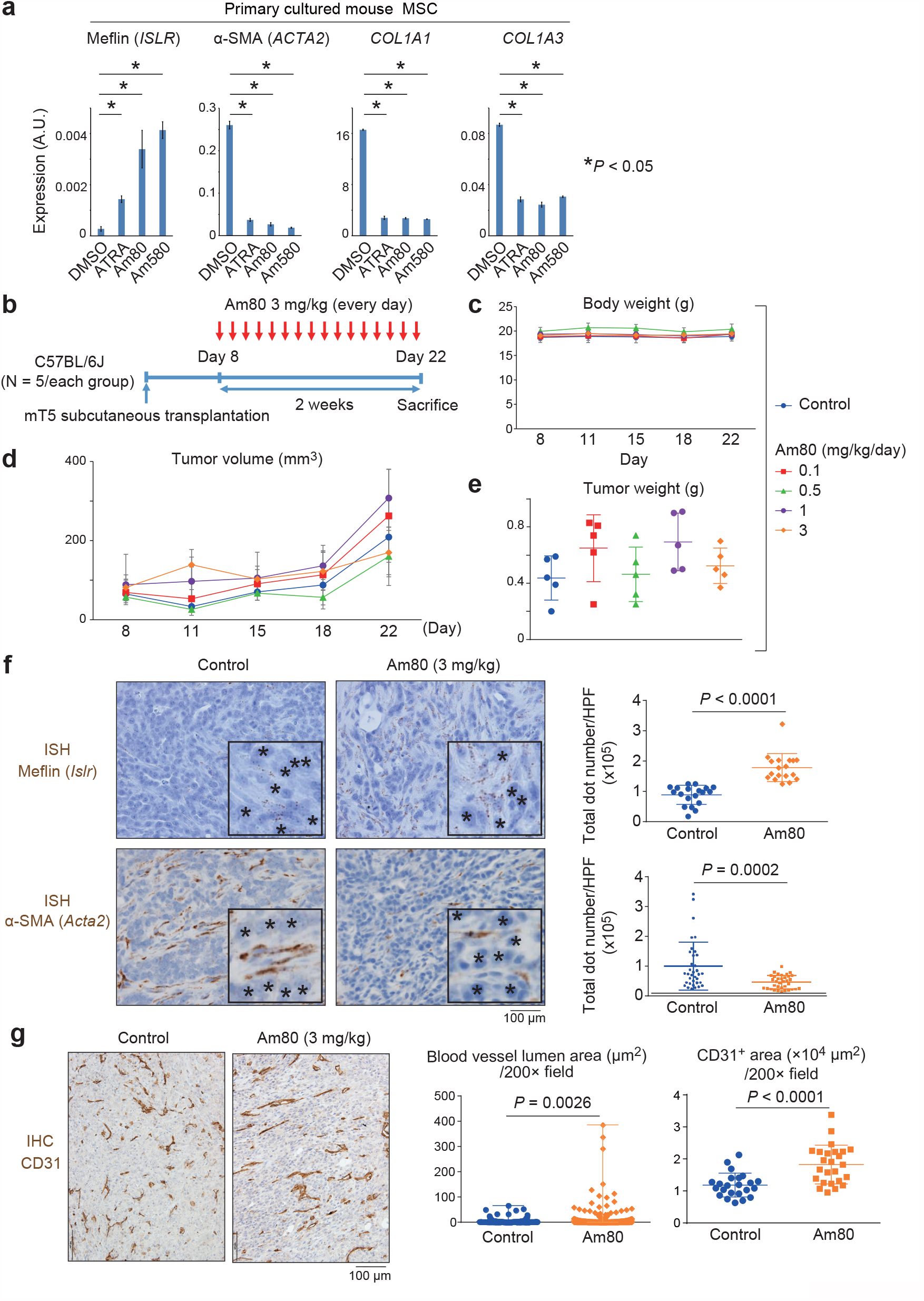
Effects of Am80 administration on the expression of Meflin in CAFs and on tumor vessel area in a PDAC xenograft mouse model. **(a)** Primary cultured mouse MSCs were plated in the presence of DMSO, ATRA, Am80, and AM580 for 48 h, and qPCR of the indicated genes was performed. **(b–e)** WT female mice (P42) were subcutaneously transplanted with mT5 mouse PDAC cells (1 × 10^6^ cells/mouse), followed by oral administration of Am80 (3 mg/kg, 0.5% carboxymethylcellulose [CMC] solution) every day for 14 days **(b)**. Body weights of mice **(c)** and tumor volumes **(d)** were measured during the observation period. **(e)** Tumor weight on the day of sacrifice (day 22). **(f, g)** Tissue sections prepared from control mT5 tumors (left) and those administered Am80 (right) were examined for Meflin and α-SMA expression by ISH **(f)** and for CD31 expression by IHC **(g)**. Meflin and α-SMA expression levels **(f)** and tumor vessel lumen and CD31^+^ areas **(g)** were calculated. For quantification of ISH signals, the number of dots per high-power (400×) microscopic field (HPF) was counted, and 20 HPFs randomly selected from 5 animals were evaluated for each group. Asterisks denote tumor cells that were negative for Meflin and α-SMA expression.

Oral administration of varying doses of Am80 to WT mice harboring subcutaneous mT5 tumors affected neither body weight nor tumor volume and weight (**Fig. 3b–e**). Am80 also did not affect the proliferation of cultured mT5 PDAC cells at concentrations of 0.01–1 µM in culture, indicating that Am80 did not have direct cytotoxic effects in mT5 PDAC cells (**Supplementary Fig. 4**). ISH analysis of mT5 tumor tissues showed that Meflin expression in stromal cells was highly upregulated by Am80 administration, accompanied by downregulation of α-SMA (**Fig. 3f**). As expected from analyses of human PDAC biopsy samples and mT5 tumors transduced with SeV-Meflin (**Figs. 1, 2**), Am80 administration increased tumor vessel area in the mT5 tumor transplantation model (**Fig. 3g**).

### Am80 administration enhanced the effects of chemotherapeutics

We next orally administered Am80 to WT mice harboring mT5 tumors for 1 week, followed by treatment with Am80 plus Gem (**Fig. 4a**). The combined treatment did not affect the body weight of mice (**Fig. 4b**). Interestingly, Am80 administration enhanced the antitumor effects of Gem, as demonstrated by monitoring of tumor volume and weight (**Fig. 4c, d**). Histological analysis using hematoxylin & eosin (H&E) staining showed that tumors treated with the combination therapy contained a greater number of necrotic cells with pyknotic nuclei (**Fig. 4e**) and exhibited higher Meflin and lower α-SMA expression than tumors treated with Gem alone (**Fig. 4f–g**). Consistent with the effects of Am80 administration on tumor vessel area (**Fig. 3g**), the concentration of Gem (2’,2-difluorodeoxycytidine [dFdC]) was higher in tumors treated with the combination therapy than in those treated with Gem monotherapy, suggesting that Am80 enhances the intratumoral delivery of Gem (**Fig. 4h**).

**Figure 4.**
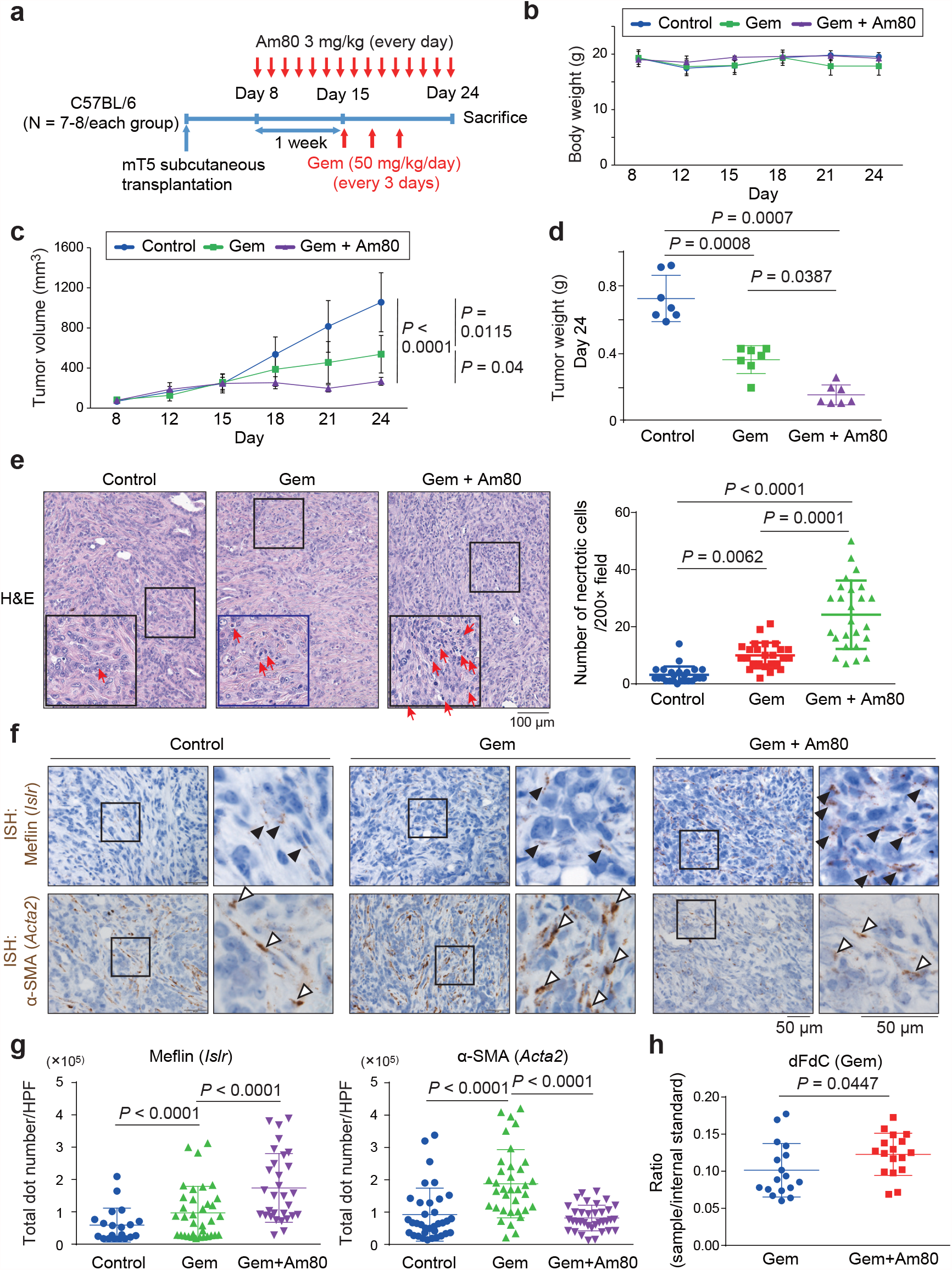
Am80 administration improved tumor sensitivity to Gem and induced Meflin expression in CAFs in a PDAC xenograft mouse model. **(a)** WT female mice (P42) were subcutaneously transplanted with mT5 mouse PDAC cells (1 × 10^6^ cells/mouse) on day 0, followed by oral administration of Am80 from day 8 for 16 consecutive days. The mice were administered Gem from day 15 three times every 3 days and sacrificed on day 24. **(b–d)** Measurement of the body weights of mice **(b)** and tumor volumes **(c)** during the observed period and of tumor weights on day 24 **(d)**. **(e)** Representative images of H&E-stained tissue sections of tumors prepared from the indicated groups. Boxed regions are magnified in insets. Arrows denote necrotic cells with pyknotic nuclei. The numbers of necrotic cells were counted and quantified in the graph shown in the right panel. Twenty-five images obtained from five tumors per group using a 20× objective lens were analyzed and quantified. **(f, g)** Tissue sections of tumors from the indicated groups were stained for Meflin and α-SMA by ISH **(f)**, followed by quantification of total dot numbers **(g)**. Open and solid arrowheads denote Meflin^+^ and α-SMA^+^ CAFs, respectively. **(h)** Quantification of dFdC (Gem) delivered to tumors in the indicated groups after administration of Gem three times. Lysates prepared from mT5 tumors treated with Gem (n = 17) and Gem plus Am80 (n =17) were added with stable isotope-labeled dFdC (internal control) subjected to liquid chromatography-mass spectrometry analysis, followed by quantification of dFdC and the internal control. The data are expressed as relative ratios of dFdC to the internal control.

The effects of oral Am80 administration were further tested using the KPC PDAC mouse model (Kras^LSL-G12D/+^; Trp53^LSL-R172H/+^; Pdx-1-Cre), which recapitulates human PDAC with extensive stromal fibroinflammatory reaction (27) and a human PDAC xenograft model (**Supplementary Fig. 5**). Echographic measurement of tumor sizes in KPC mice revealed that Am80 administration significantly improved the antitumor effects of Gem (**Supplementary Fig. 5a–c**). We next transplanted BxPC3 human PDAC cells into immunocompromised nude mice, followed by oral administration of Am80 and the combination chemotherapy of Gem and nabPTX (**Supplementary Fig. 5d**). Am80 administration did not alter the body weight of the mice but significantly improved the antitumor effects of Gem plus nabPTX (**Supplementary Fig. 5e, f**). This observation suggested minor involvement of the immune machinery in the mechanism of action.

### Am80-mediated enhancement of chemosensitivity required Meflin expression in CAFs

Next, to further confirm the significance of Am80-mediated Meflin expression in enhanced chemosensitivity, we transplanted mT5 PDAC cells into Meflin-KO mice, followed by Am80 administration and Gem treatment (**Fig. 5a**). Gem treatment, with or without Am80 administration, did not alter the body weight of the transplanted Meflin-KO mice (**Fig. 5b**). Am80 administration in combination with Gem resulted in a slight but not statistically significant decrease in tumor volume and weight compared with Gem monotherapy (**Fig. 5c, d**). The numbers of necrotic cells with pyknotic nuclei and the expression levels of α-SMA in the stroma were comparable between the monotherapy and combination therapy groups (**Fig. 5e–g**). Because Meflin is specifically expressed in CAFs but not other cell types in PDAC tissue (20), these data suggested that Am80 exerted its effects on chemotherapeutic efficacy by upregulation of Meflin in CAFs. Indeed, when we transplanted BxPC3 PDAC cells with or without immortalized human PSCs into nude mice, followed by combination therapy with Am80 and Gem (**Fig. 5h**), we found that effects of Am80 administration on tumor sensitivity to Gem were observed only when BxPC3 cells were cotransplanted with PSCs (**Fig. 5i, j**). Thus, enhancement of tumor chemosensitivity by Am80 was mediated by its effects on CAFs but not cancer cells.

**Figure 5.**
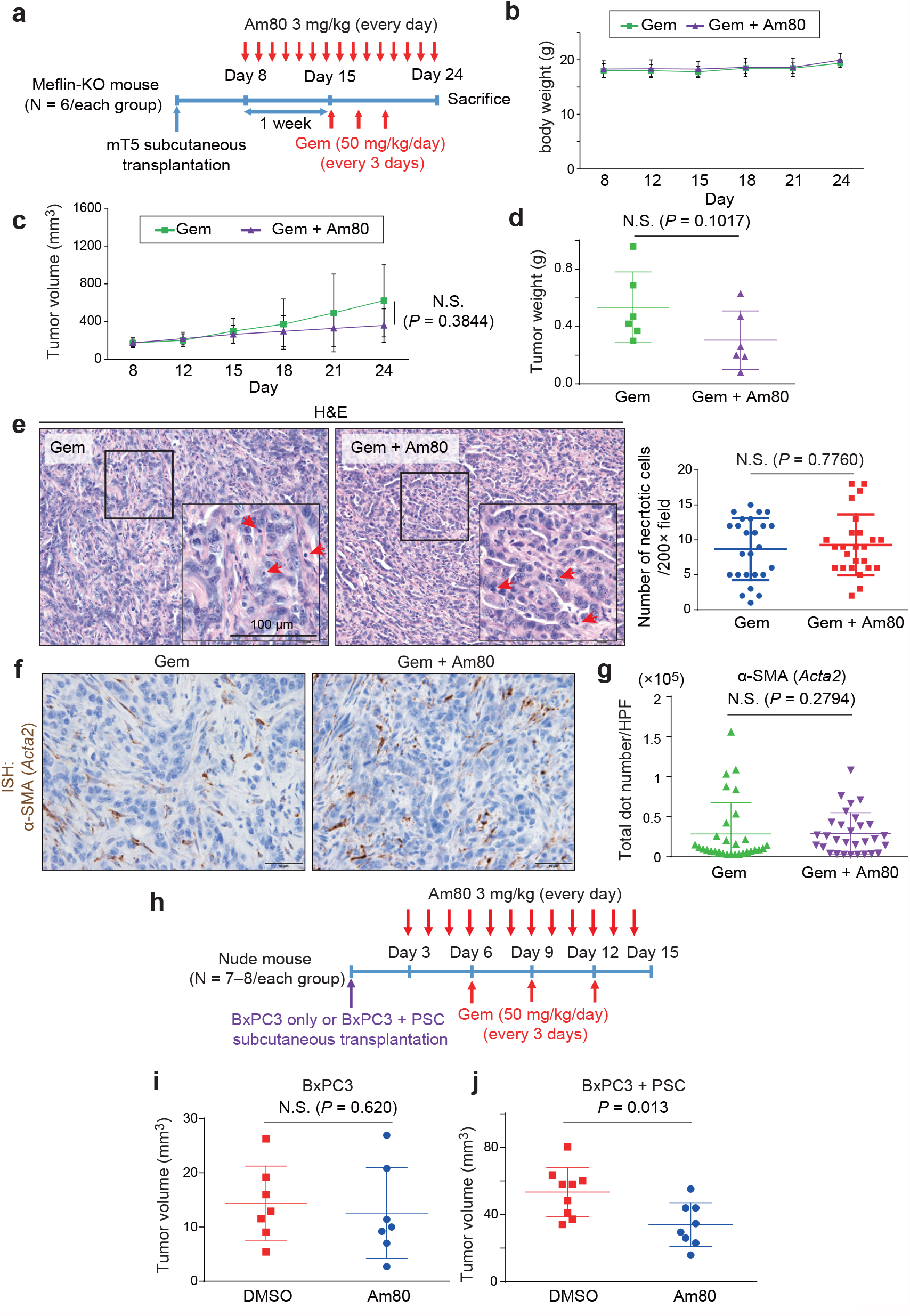
Increased tumor chemosensitivity by Am80 administration was mediated by alterations in Meflin expression in CAFs. **(a)** Meflin-KO female mice (P42) were subcutaneously transplanted with mT5 mouse PDAC cells (1 × 10^6^ cells/mouse) on day 0, followed by oral administration of Am80 from day 8 for 16 consecutive days. Mice were administered Gem from day 15 three times every 3 days and sacrificed on day 24. **(b–d)** Measurement of the body weights of mice **(b)** and tumor volumes **(c)** during the observation period, and tumor weights on day 24 **(d)**. **(e)** Representative images of H&E-stained tissue sections of tumors prepared from the indicated groups. Boxed regions are magnified in insets. Arrows denote necrotic cells with pyknotic nuclei. The numbers of necrotic cells were counted and quantified in the graph shown in the right panel. **(f, g)** Tissue sections of tumors from the indicated groups were stained for α-SMA by ISH **(f)**, followed by quantification of the total dot numbers **(g)**. **(h–j)** Adult nude female mice (P42) were subcutaneously implanted with either human BxPC3 PDAC cells alone (1 × 10^6^ cells/mouse) or both BxPC3 cells and immortalized human PSC cells at a 1:5 ratio (BxPC3: 1 × 10^6^ cells, PSC: 5 × 10^6^ cells), followed by oral administration of either DMSO or Am80 from day 3 for 12 consecutive days **(h)**. Mice were administered Gem from day 6 three times every 3 days, followed by sacrifice on day 15 and measurement of tumor volumes **(i, j)**.

### Meflin interacted with lysyl oxidase (Lox) to suppress its collagen crosslinking activity

Meflin augments bone morphogenetic protein (BMP) 7 signaling to oppose the profibrotic action of transforming growth factor (TGF)-β (36, 37-); however, this mechanism does not fully explain the changes found in tumors administered Am80 and the differences between WT and Meflin-KO tumors. In our previous reports, second-harmonic generation (SHG) microscopic observations revealed that the stroma of tumors in Meflin-KO KPC mice exhibited significant remodeling of collagen architectures compared with those in WT KPC mice (20). These data suggested that Meflin was involved in the regulation of collagen configuration in the tumor stroma, which may subsequently change the mechanical properties of the tumor stroma, such as interstitial pressure (38).

To identify the mechanisms through which Meflin is involved in regulation of stromal collagen, we attempted to identify novel ligands of Meflin. To this end, we transduced mouse MSCs with Meflin cDNA tagged with the hemagglutinin (HA) epitope, followed by immunoprecipitation (IP) with anti-HA antibodies and analysis of the eluate by mass spectrometry (**Supplementary Fig. 6a, b**). We identified seven secreted or membrane proteins as candidate Meflin interactors and focused on the protein with the highest score, Lox (**Supplementary Fig. 6c**). Lox is known to crosslink collagen and elastin, the two major components of the ECM, contributing to the stiffness and interstitial pressure of the tumor stroma (39, 40). IP assays confirmed that Meflin interacted with Lox (**Fig. 6a**) and lysyl oxidase like 2 (Loxl2), another member of the Lox gene family (**Supplementary Fig. 6d**). Direct interaction of Meflin and Lox was demonstrated in *in vitro* binding assays using purified recombinant Meflin and Lox proteins (**Supplementary Fig. 6e**). Additional IP assays revealed the interaction of endogenous Meflin with Lox and Loxl2 proteins, highlighting the physiological relevance of these protein interactions (**Fig. 6b, Supplementary Fig. 6f**). A mutant Lox lacking the amino-terminal pro-sequence also interacted with Meflin, suggesting that the catalytic domain (CD) of Lox was responsible for this interaction (**Fig. 6c, d**). Pull-down assays using Meflin mutants fused with the constant region (Fc) of human immunoglobulin lacking one of the leucine-rich repeats (LRRs) and the immunoglobulin (Ig)-like domain showed that a region spanning the LRR1–4 domains of Meflin was likely to be responsible for Lox binding (**Supplementary Fig. 7a, b**). Functionally, biochemical assays showed that Meflin had inhibitory effects on the oxidative deamination activity of Loxl2, suggesting that Meflin had inhibitory roles in collagen crosslinking mediated by Lox or Loxl2 (**Fig. 6e**).

**Figure 6.**
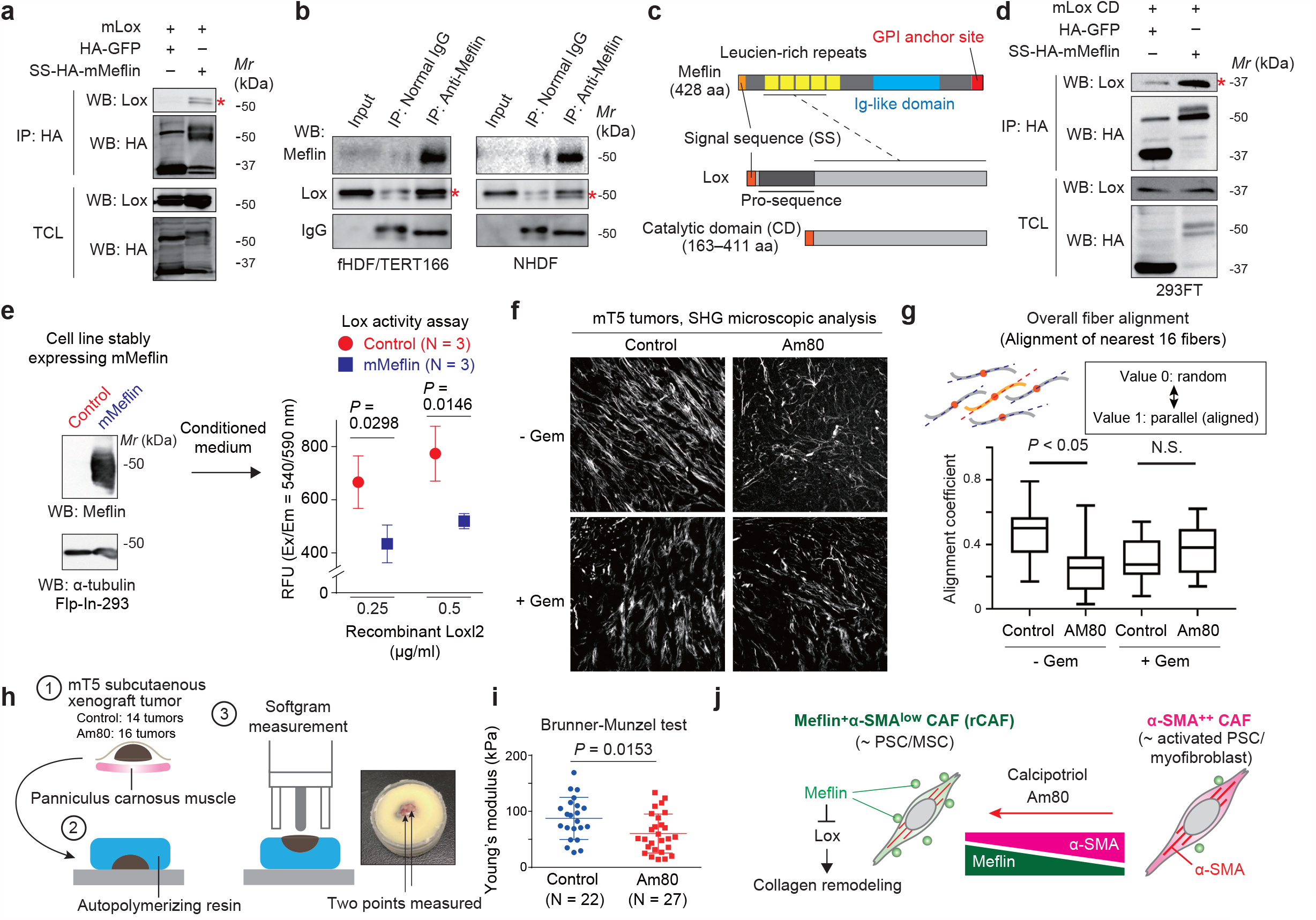
Meflin was involved in the regulation of Lox activity and collagen remodeling. **(a)** The indicated plasmids were transfected into HEK293 cells, followed by IP with anti-HA antibodis and WB analysis. Lox coprecipitated with HA-Meflin (asterisk). mMeflin, mouse Meflin; mLox, mouse Lox; SS, signal sequence; TCL, total cell lysate. **(b)** Interaction of endogenous Meflin and Lox proteins. Cell lysates prepared from the fHDF/TERT166 fibroblast cell line (left) or primary cultured human fibroblasts (NHDFs, right) were immunoprecipitated with anti-Meflin antibodies, followed by analysis of eluates by WB with the indicated antibodies. Lox proteins coprecipitated with Meflin (asterisk). **(c)** Primary structure of human Meflin (top) and mouse Lox (middle). Lox comprises an amino-terminal SS, a pro-sequence, and a catalytic domain (CD). (Bottom) Construct encoding the CD of Lox with an SS at the amino terminus. Dashed line: protein domains responsible for the Lox/Meflin interaction. **(d)** The indicated plasmids were cotransfected into 293FT cells, followed by IP with anti-HA antibodies and WB analysis. Meflin interacted with the CD of mLox (asterisk). **(e)** Conditioned medium from control Flp-In-293 cells or Flp-In-293 cells stably expressing mMeflin was mixed with recombinant Loxl2 (0.25 and 0.5 µg/mL), followed by measurement of hydrogen peroxide using a fluorometric-based method. (Left) WB analysis of Meflin expression in Flp-In-293 cells. RFU, relative fluorescence unit. **(f, g)** Measurement of collagen alignment in the stroma of mT5 tumors developed in mice by SHG microscopy. Eight to sixteen images from tissue sections of control tumors and tumors treated with Am80 were analyzed. Representative images **(f)** and quantification of collagen alignment **(g)**. **(h, i)** Measurement of the stiffness of mT5 tumors developed in control mice (n = 14) and those treated with Am80 (n = 16). The Young’s modulus of the flat surfaces of tumors contacting the fascia of the panniculus carnosus muscle were measured (**h**). One or two points were selected in each tumor, and each point was measured more than 5 times; quantification was performed (**i**). **(j)** Schematic diagram showing the working hypothesis for the mechanism of Am80-mediated alteration in tumor sensitivity to chemotherapeutics.

We next analyzed the effects of Am80 administration on the collagen signature of mT5 xenograft tumors in the presence or absence of Gem by SHG microscopy (**Fig. 6f, g**). The data showed highly aligned collagen fibers in control mT5 tumors, which were significantly reversed by Am80 administration, in the absence of Gem treatment. This difference disappeared upon Gem treatment. Other collagen signatures, such as curvature, width, and fiber length, were all comparable between the tested groups (**Supplementary Fig. 8a–c**). Interestingly, measurement of the Young’s modulus of the tumors showed that Am80-treated tumors were softer than control tumors (**Fig. 6h, i**). Because collagen fiber alignment modulates tumor stiffness (38), these data suggested that Am80-mediated Meflin upregulation inhibited Lox-mediated collagen crosslinking, thereby controlling the physical properties of tumors (**Fig. 6j**).

### Differential effects of Am80 and vitamin D administration on CAF gene expression and tumor progression

Administration of the vitamin D analogue calcipotriol alters the gene expression of stromal cells of PDAC in mice (30). Therefore, we next investigated how Am80 differed from vitamin D analogues in terms of effects on gene expression in PDAC CAFs. Thus, we stimulated primary cultured PSCs isolated from three patients with PDAC (PSC163, PSC52, and PSC119) with dimethyl sulfoxide (DMSO, control), Am80, or calcipotriol. Gene expression profiles were then compared using microarray-based transcriptomic analysis (**Supplementary Fig. 9a, b**). Extracting genes that were differentially expressed between Am80- and calcipotriol-treated cells revealed that Meflin expression was more significantly upregulated by Am80 than calcipotriol. Interestingly, the expression levels of fibroblast activation protein (FAP) and chemokine (C-C motif) ligand 2 (CCL2), which are markers of pCAFs (2, 6, 7, 41, 42-), were higher in Am80-treated PSCs than calcipotriol-treated PSCs, whereas the expression levels of other pCAF marker genes, such as C-X-C motif chemokine ligand 12 (CXCL12), podoplanin (PDPN) and periostin (POSTN), were higher in calcipotriol-treated cells. These data suggested that Am80 and calcipotriol differentially regulated gene expression in PSCs.

We finally treated mT5 xenograft tumors with Gem in combination with Am80 or calcipotriol administered orally or intraperitoneally (i.p.), respectively (**Supplementary Fig. 9c**). Both treatments did not result in decreased body weight (**Supplementary Fig. 9d**). Interestingly, administration of Am80 but not calcipotriol significantly improved Gem efficacy in the xenograft model (**Supplementary Fig. 9e**). These results, although not conclusive and possibly confounded by different doses and timings of Am80 and calcipotriol administration, suggested that Am80 and calcipotriol had distinct effects on gene expression in CAFs, which may affect the efficacy of chemotherapeutics.

## Discussion

In the current study, we found that the number of Meflin^+^ CAFs, a candidate for rCAFs in PDAC, correlated with the response of patients with PDAC to chemotherapies. We also demonstrated the tumor-suppressive role of Meflin in PDAC mouse models by inducing its expression through genetic and pharmacological approaches. These experiments led to the identification of Am80 as an inducer of Meflin expression in CAFs and showed that Am80 improved the sensitivity of PDAC to chemotherapeutics. Finally, we showed that Meflin functioned to suppress Lox collagen crosslinking activity, which may increase tumor vessel area and chemotherapeutic sensitivity, as observed in tumors with high Meflin expression (**Fig. 6j**). Taken together, these data support that there are functionally heterogeneous but phenotypically plastic subsets of CAFs in the tumor stroma, consistent with the observation that tumor immunity is regulated by the net balance of anti- and protumor immune cells (43, 44). Given that Meflin^+^ CAFs weakly or moderately express α-SMA (7, 20), we speculate that the failure to deplete all α-SMA^+^ CAFs in a preclinical PDAC model was caused by the depletion of Meflin^+^ CAFs (10).

The findings in the current study should be discussed in the context of a previous study, in which researchers demonstrated that calcipotriol administration suppresses stromal activation and improves the efficacy of Gem treatment in a PDAC mouse model (30). Calcipotriol targets activated PSCs expressing vitamin D receptor (VDR), which likely represent pCAFs, to revert or deactivate them to a more quiescent, less cancer-supportive state (20, 30). This VDR and ligand-mediated reversion of activated PSCs to quiescent PSCs has been called stromal reprogramming, which is the basis of ongoing clinical trials that test the efficacy of vitamin D analogs in combination with chemotherapeutics or immune checkpoint inhibitors in patients with PDAC (www.clinicaltrials.gov). We previously reported that Meflin is a marker of quiescent PSCs in the normal pancreas, and its expression was upregulated by calcipotriol (20). This was consistent with the results of a present study showing that Meflin expression in CAFs correlates with sensitivity to chemotherapeutics in PDAC. Therefore, Meflin may be a surrogate or monitoring marker of the stromal reprogramming of PDAC in clinical trials.

Interestingly, our experiments showed that Am80 was a more potent inducer of high chemosensitivity than calcipotriol in a xenograft tumor model. However, these findings may be limited to the experimental models used in the current study, and further studies on more human-relevant models and clinical trials are warranted. Am80 is a synthetic unnatural retinoid that specifically binds to RARα and β, with Ki values of 6.5 × 10^−9^ and 3.0 × 10^−10^ M, respectively, but not RAR γ; these findings are contradictory to ATRA, which has similar affinities to all RAR isoforms (45). In addition, Am80 shows several pharmacological advantages over ATRA. For example, Am80 exhibits significantly higher stability to light, heat, and oxidation than ATRA, which allows a lower dose (6 mg/m^2^) to exert its effect than ATRA (45 mg/m^2^) in clinical practice (35). Am80 binds to cellular RA-binding protein, the induced upregulation of which is known to mediate resistance to ATRA in APL, with a 20-fold lower affinity than ATRA (35, 45). Oral intake of Am80 does not produce serious metabolic complications, such as hypercalcemia caused by high-dose vitamin D administration (46). Furthermore, similar to ATRA, Am80 has been shown to have various biological activities, including induction of cell differentiation and modulation of immunity (33, 47). Therefore, the hypothesis that Am80 suppresses tumor progression by increasing the number of Meflin^+^ rCAFs may be a simplistic interpretation of the experimental results obtained in the current study, and more extensive studies are needed to comprehensively elucidate the effects of Am80 on tumor cells and TME components. Additionally, it will be necessary to evaluate the effects of Am80 oral administration on drug sensitivity in patients with PDAC in the clinical settings.

Our data showed a significant increase in blood vessel areas in tumors treated with SeV-Meflin and Am80, supporting the involvement of these interventions in modulation of tumor angiogenesis. However, previous studies showed an inhibitory effect of Am80 on angiogenesis (48) and no Meflin expression in vascular endothelial and smooth muscle cells (21); thus, it is plausible that SeV-Meflin- and Am80-mediated changes in the tumor vessel lumen may be attributed to alterations in the mechanical properties of the tumor stroma. Consistent with this hypothesis, our search for Meflin ligands revealed that Meflin interacts with Lox to inhibit its collagen crosslinking activity. Lox is a critical regulator of the physical properties of the tumor stroma, such as stiffness and interstitial pressure (38). Therefore, artificial induction of Meflin expression in CAFs may be a rational approach to improve the perfusion of tumor vessels and drug delivery. The involvement of other reported functions of Meflin, including the regulation of BMP7, Wnt, and Hippo-Yes-associated protein (Yap) signaling pathways, in the regulation of tumor vessel area also needs to be considered (49, 50).

In conclusion, our current findings showed that Meflin expression in CAFs improved tumor sensitivity to chemotherapeutics, consistent with our previous studies identifying Meflin as a marker of rCAFs, which are functionally different from protumorigenic pCAFs (7, 22, 43). Consistent with this, the induction of Meflin expression by either genetic or pharmacological approaches improved tumor sensitivity to chemotherapeutics in a PDAC xenograft model. Taken together, these data provide a rationale for therapeutic strategies that increase the number of rCAFs in the treatment of patients with PDAC.

## Acknowledgments

We thank David Tuveson (Cold Spring Harbor Laboratory) and Chang-il Hwang (UC Davis College of Biological Sciences) for providing the mouse PDAC cell line mT5; Kohji Kusano (ID Pharma Co., Ltd.) for generating recombinant Sendai virus; Shuzo Watanabe, Kaoru Shimada (RaQualia Pharma Inc.), and Hisao Ekimoto (TMRC Co., Ltd.) for helpful discussions on Am80; Kentaro Taki (Nagoya University) for help with mass spectrometry; and Kozo Uchiyama and Kaori Ushida (Nagoya University) for technical assistance. This work was supported by a Grant-in-Aid for Scientific Research (B) (grant nos. 18H02638 to A.E. and 20H03467 to M.T.) commissioned by the Ministry of Education, Culture, Sports, Science and Technology of Japan; Nagoya University Hospital Funding for Clinical Research (to A.E.); AMED-CREST (Japan Agency for Medical Research and Development, Core Research for Evolutional Science and Technology; grant nos. 20gm0810007h0105 and 20gm1210009s0102 to A.E.); and the Project for Cancer Research and Therapeutic Evolution (P-CREATE) from AMED (grant nos. 20cm0106377h0001 to A.E. and 21cm0106704h0002 to Y.M.).

## Author contributions

T.Ii. designed and performed the experiments, analyzed the data, and wrote the manuscript. Y.M. and N.E. designed and performed the experiments and analyzed the data. S.M.P. and B.M.B. performed SHG analysis. L.W. performed a mass spectrometric analysis. K.Ku. performed the measurement of intratumoral concentrations of dFdC. K.Ka., A.M., T.Is., E.O., and H.K. provided the clinical samples and intellectual input. S.I. and H.H. assisted with the analysis of tumor stiffness. S.M. and Y.S. assisted with histological analysis. H.K., Y.H., M.F., and M.T. directed the project and provided intellectual input. A.E. directed the project and wrote the manuscript.

