## Supplementary Data for "Conversion of cancer-associated fibroblasts from pro- to antitumor improves the sensitivity of pancreatic cancer to chemotherapeutics"

Iida *et al.*

**Supplementary Data**

**1. Supplementary Figures and legends**

**2. Supplementary Methods**

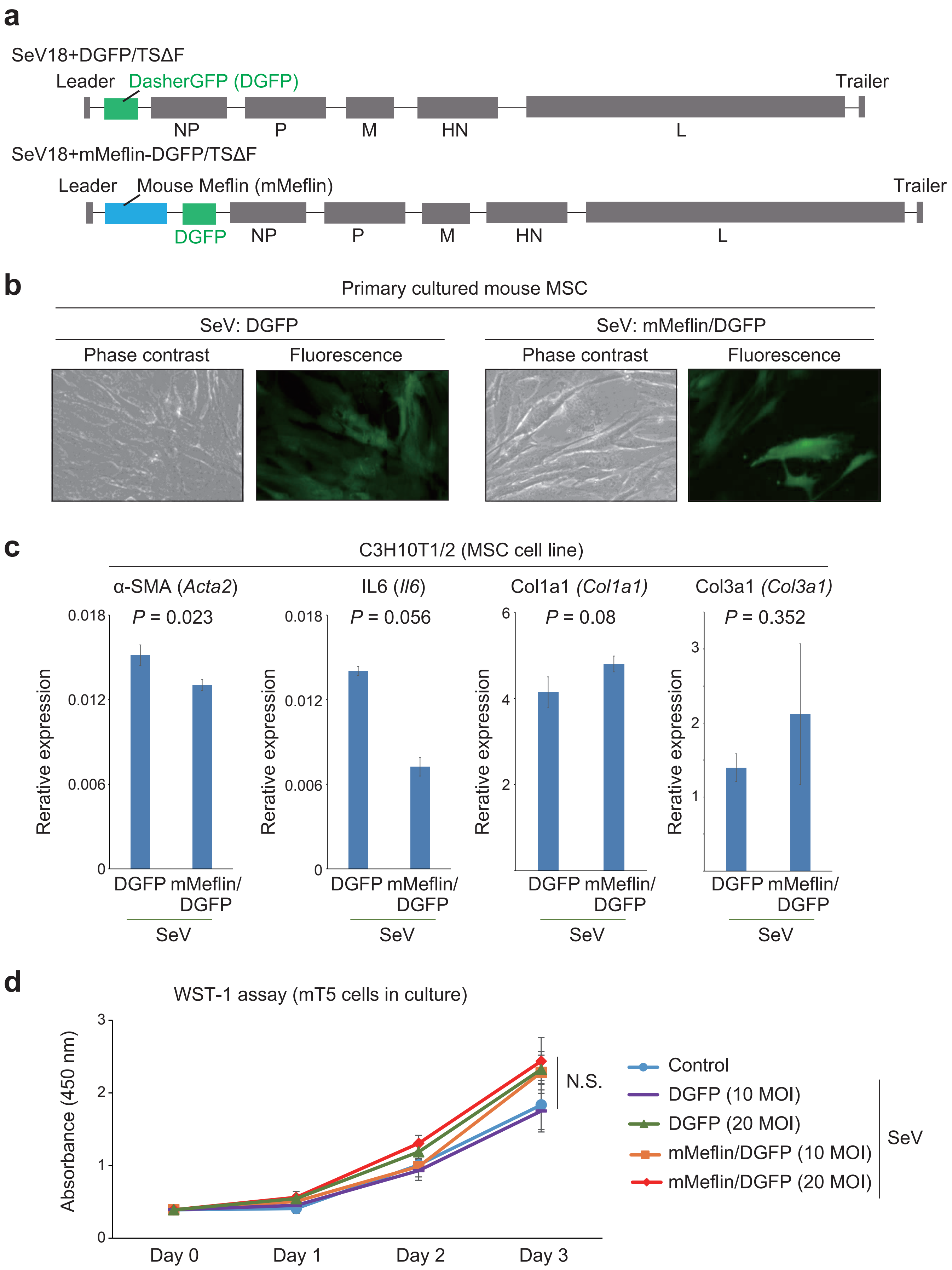


**Supplementary Fig 1**

**Supplementary Fig 1.** Effects of infection of recombinant SeV expressing Meflin on gene expression in MSCs and proliferation of mT5 PDAC cells.

**(a)** Structures of SeV vectors encoding DasherGFP (DGFP, top) and mMeflin and DGFP (bottom). The DGFP and mMeflin/DGFP cDNAs and other viral genes (NP, P, M, HN, and L) were flanked by 5' and 3' extracistronic regions known as the leader and trailer, respectively.

**(b)** Phase contrast and fluorescent images of primary cultured mouse MSCs infected with recombinant SeV-DGFP (left) and SeV-mMeflin/DGFP (right).

**(c)** Recombinant SeV-DGFP or SeV-mMeflin/DGFP were used to infect mouse MSCs (C3H10T1/2), and qPCR of the indicated genes was performed.

**(d)** Proliferation of cultured mT5 PDAC cells infected with SeV-DGFP and SeV-mMeflin/DGFP at the indicated multiplicity of infection (MOI) was examined by WST-1 assay.

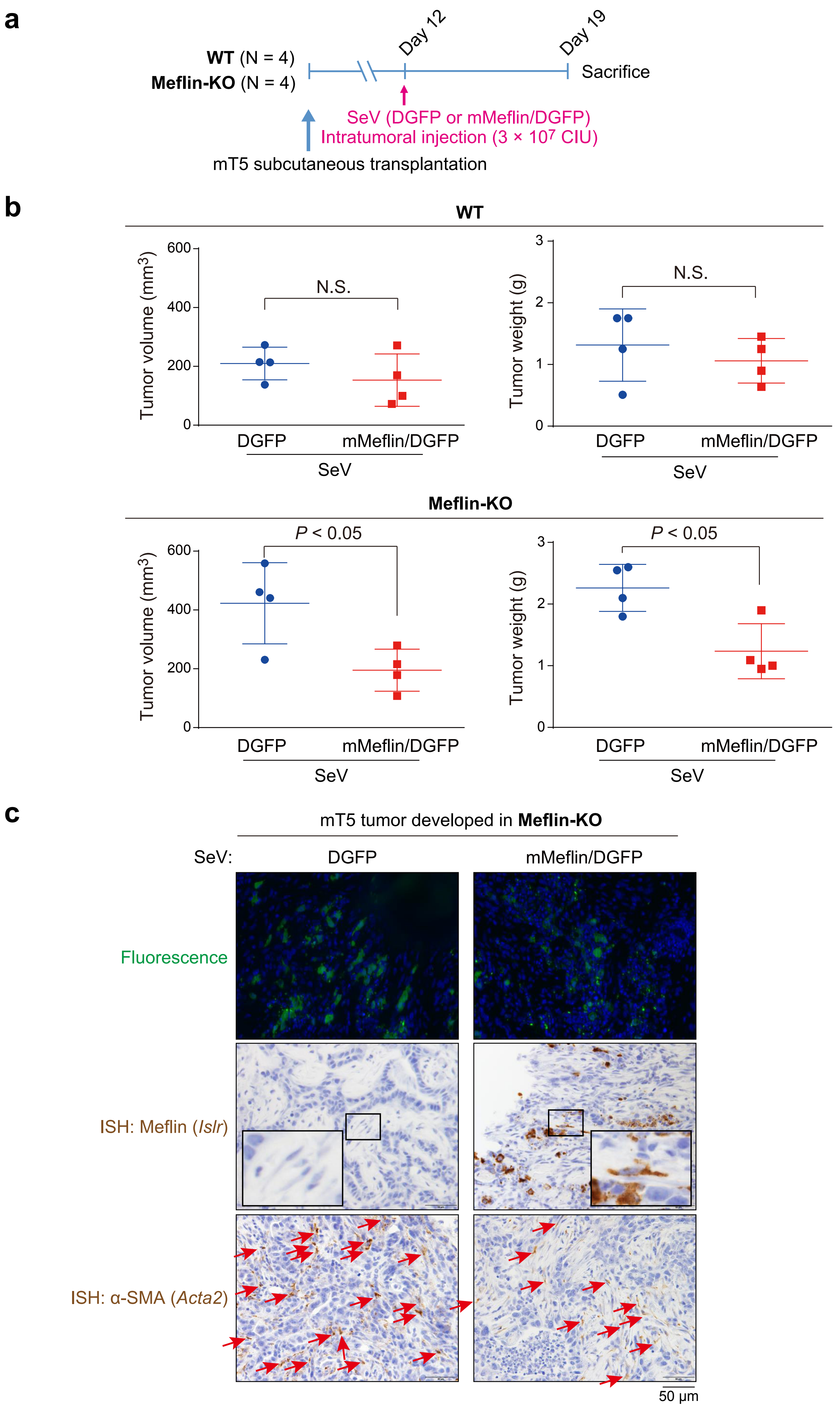


Supplementary Fig 2

**Supplementary Fig 2.** Transduction of SeV-Meflin exerted antitumor effects in Meflin-KO but not WT mice.

**(a)** WT or Meflin-KO female mice (P42) were subcutaneously transplanted with mT5 mouse PDAC cells ($1 \times 10^6$ cells/mouse), followed by intratumoral injection of SeV-DGFP and SeV-Meflin/DGFP ($3 \times 10^7$ cell CIU) on day 12. Mice were sacrificed on day 19.

**(b)** Measurement of tumor volumes (left) and weights (right) for tumors injected with SeV-DGFP or SeV-Meflin/DGFP in WT (upper panel) and Meflin-KO (lower panels) mice.

**(c)** Observation of GFP fluorescence in frozen sections prepared from mT5 tumors infected with SeV-DGFP and SeV-Meflin/DGFP (upper panel). FFPE sections were also obtained from tumors and stained for Meflin and α-SMA by ISH (middle and lower panels). Boxed regions are magnified in insets. Arrows indicate α-$SMA^+$ CAFs.

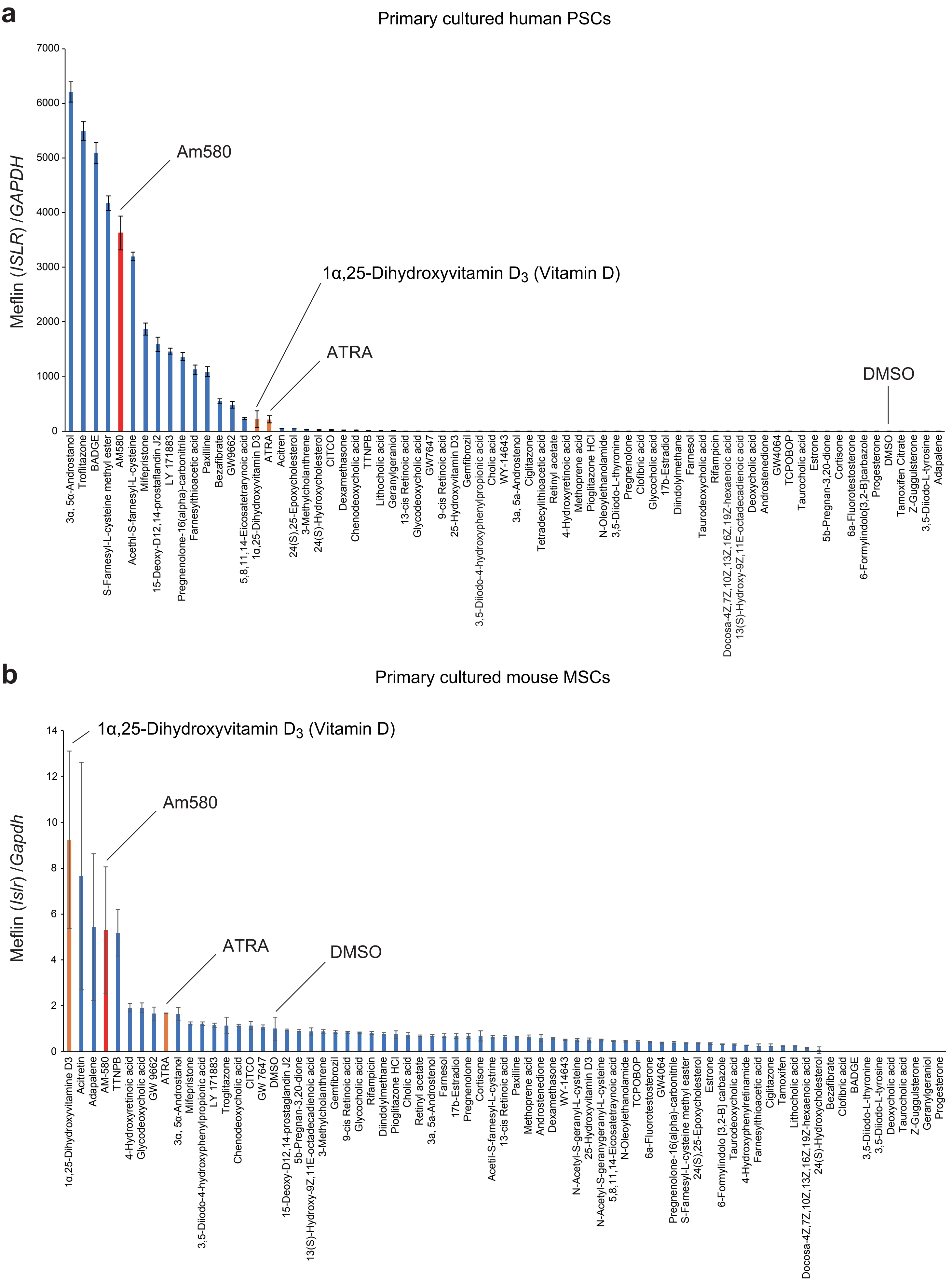


**Supplementary Fig 3**

**Supplementary Fig 3.** Screening of a library of ligands of the nuclear receptor superfamily to identify compounds upregulating Meflin in human PSCs and mouse MSCs.

**(a, b)** Primary cultured human PSCs isolated from pancreatic tissue adjacent to PDAC **(a)** and mouse MSCs **(b)** were plated in 6-well plates and allowed to reach superconfluence. The indicated compounds were then added at 1 μM, and the cells were incubated for 48 h. Meflin mRNA levels were determined by qPCR. All samples were normalized to GAPDH and expressed as fold change to the control (DMSO).

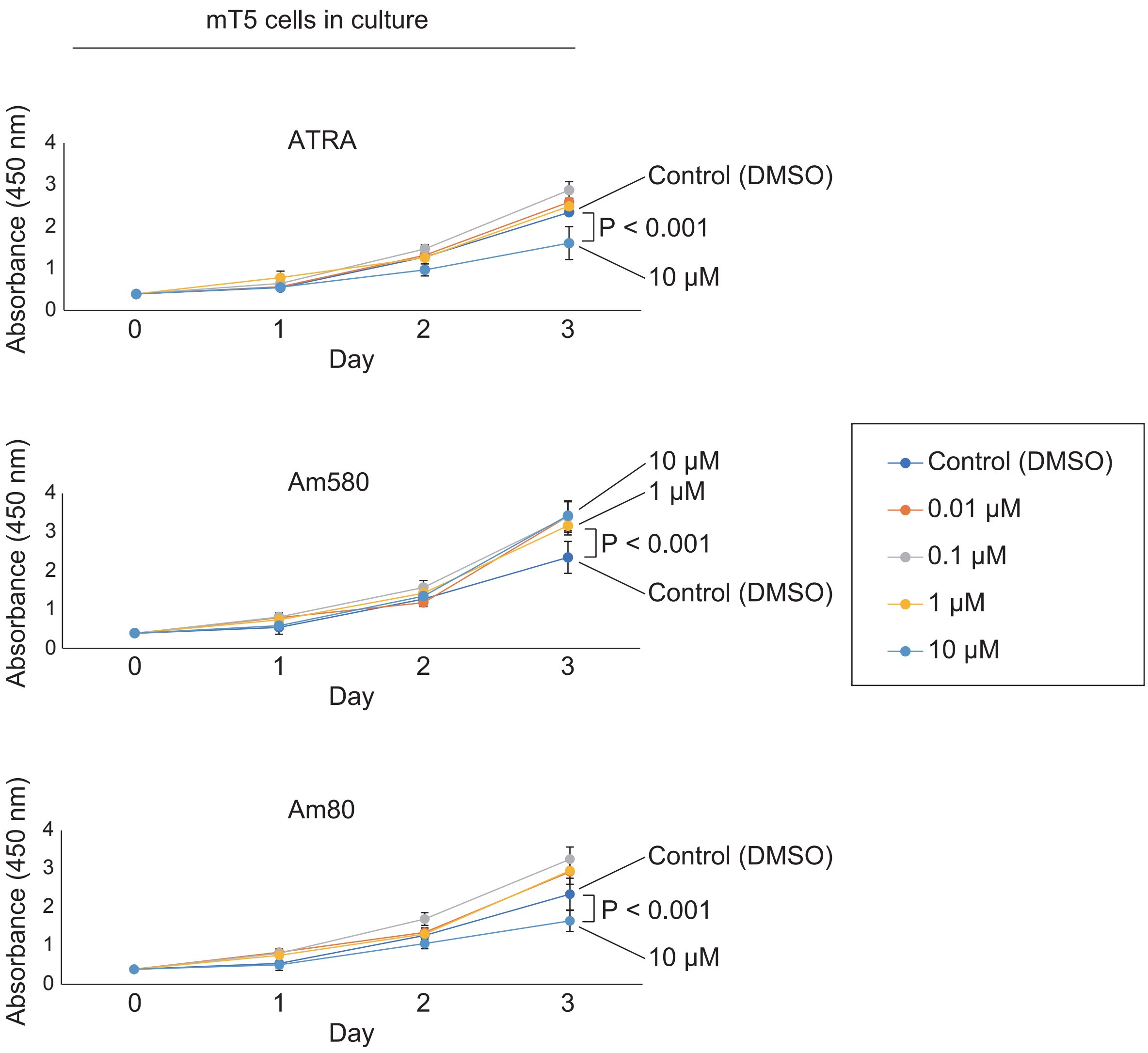


**Supplementary Fig 4**

**Supplementary Fig 4.** Minor effects of ATRA, Am580, and Am80 on the proliferation of mT5 cells.

mT5 cells ($1 \times 10^3$ cells) were plated on wells of a 96-well plate and cultured for 3 days in the presence of ATRA, Am580, and Am80 at the indicated concentrations. Cell proliferation was evaluated by WST-1 assays each day, and the data were quantified.

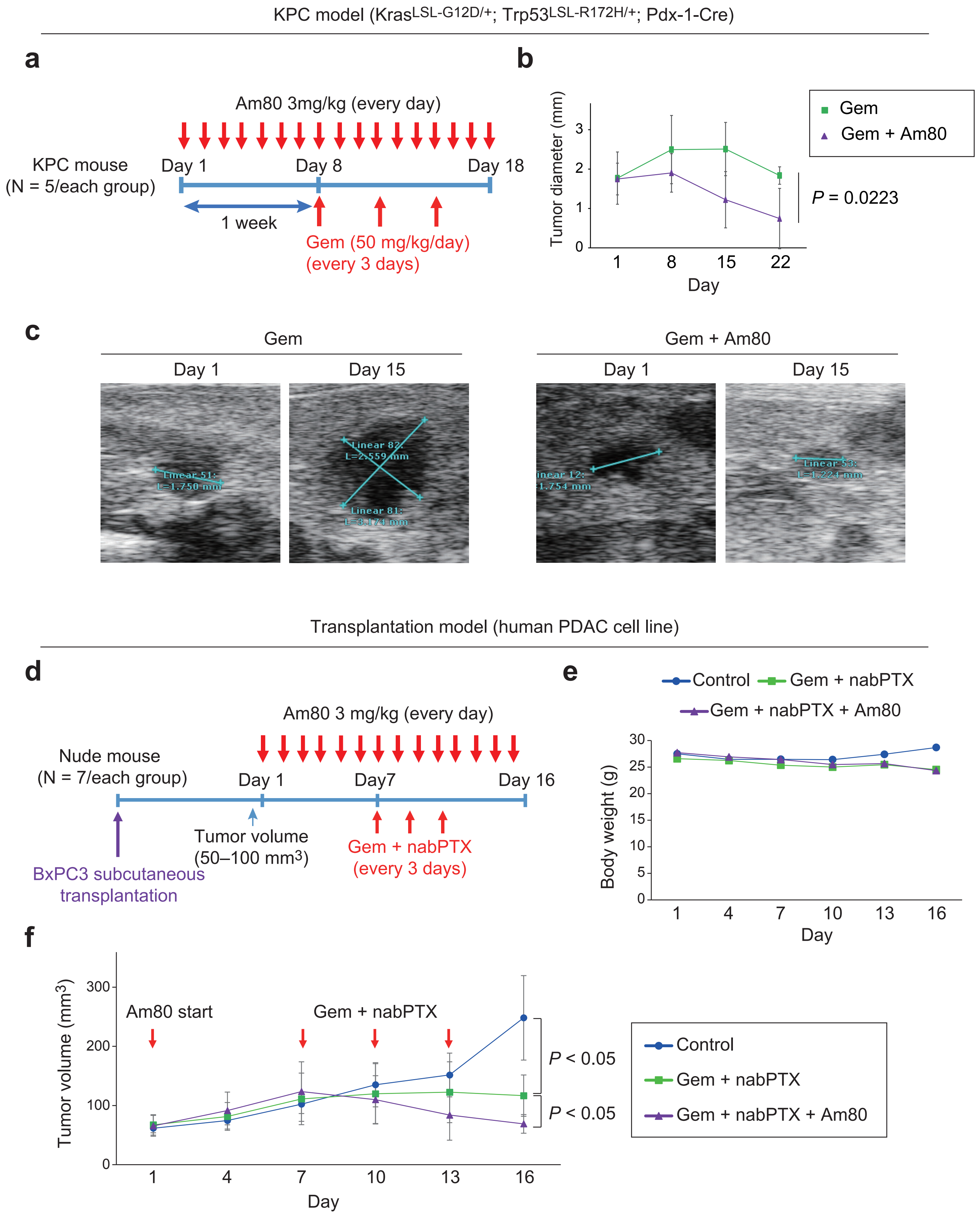


**Supplementary Fig 5**

**Supplementary Fig 5.** Effects of Am80 administration on tumor progression in the KPC PDAC mouse model and human PDAC xenografts in nude mice.

**(a)** KPC mice, which harbor mutations in the K-Ras and p53 genes in Pdx1 lineage cells, were raised until tumors in the pancreas reached approximately 1–2 mm in diameter, as measured by high-resolution ultrasound imaging (day 1). Am80 was then orally administered every day. The mice were i.p. injected with Gem three times every 3 days from day 8.

**(b)** Measurement of tumor diameters by high-resolution ultrasound imaging during the observation period.

**(c)** Representative ultrasound images of tumors treated with Gem alone (left) and the combination of Gem and Am80 (right) ont days 1 and 15.

**(d)** Adult nude female mice (P42) were subcutaneously implanted with human BxPC3 PDAC cells ($1 \times 10^6$ cells). The tumors were grown until they reached 50–100 $mm^3$ in volume, and Am80 was then orally administered for consecutive days (days 1–16). Mice were administered Gem plus nabPTX from day 7 three times every 3 days.

**(e, f)** Measurement of body weights **(e)** and tumor volumes **(f)** for the indicated groups during the observation period.

**a**

pRetroQ-SS-Flag-HA-mMeflin
pRetroQ-GFP
(SS: signal sequence)

↓ Retroviral infection

Mouse MSCs

↓

IP with HA antibody

↓

Elution with HA peptide

↓

LC/MS (Q Exactive)

**b**

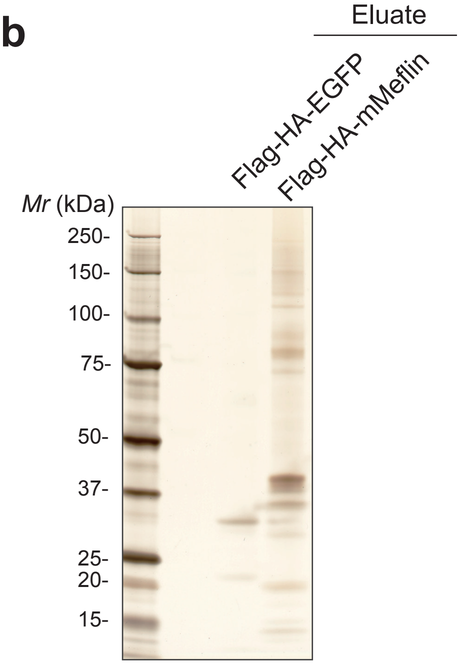


**c**

Secreted or membrane proteins detected in Meflin immunoprecipitates from mouse MSCs (Q Exactive)

| | Protein name [UniProt Knowledgebase name] | Score GFP | Score HA-mMeflin | Coverage GFP | Coverage HA-mMeflin |
|---|---|---|---|---|---|
| → | Protein-lysine 6-oxidase [LYOX_MOUSE] | - | 129.3 | - | 13.63 |
| | CD44 antigen [CD44_MOUSE] | - | 55.1 | - | 1.54 |
| | Fibrocystin-L [PKHL1_MOUSE] | - | 50.3 | - | 0.14 |
| | Bone marrow stromal antigen 2 [BST2_MOUSE] | - | 49.5 | - | 5.23 |
| * | Immunoglobulin superfamily containing leucine-rich repeat protein [ISLR_MOUSE] | - | 47.8 | - | 2.34 |
| | Lactadherin [MFGM_MOUSE] | - | 34.8 | - | 4.10 |
| | Probable G-protein coupled receptor 61 [GPR61_MOUSE] | - | 26.2 | - | 1.78 |
| | Cadherin-13 [CAD13_MOUSE] | - | 26.0 | - | 1.96 |

Cytoplasmic proteins identified by the LC/MS analysis are not listed

**d**

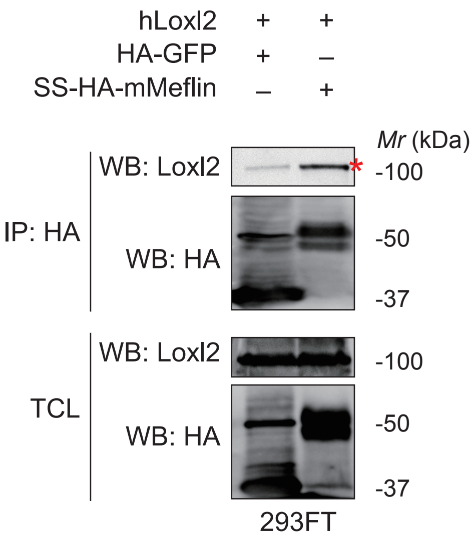


**e**

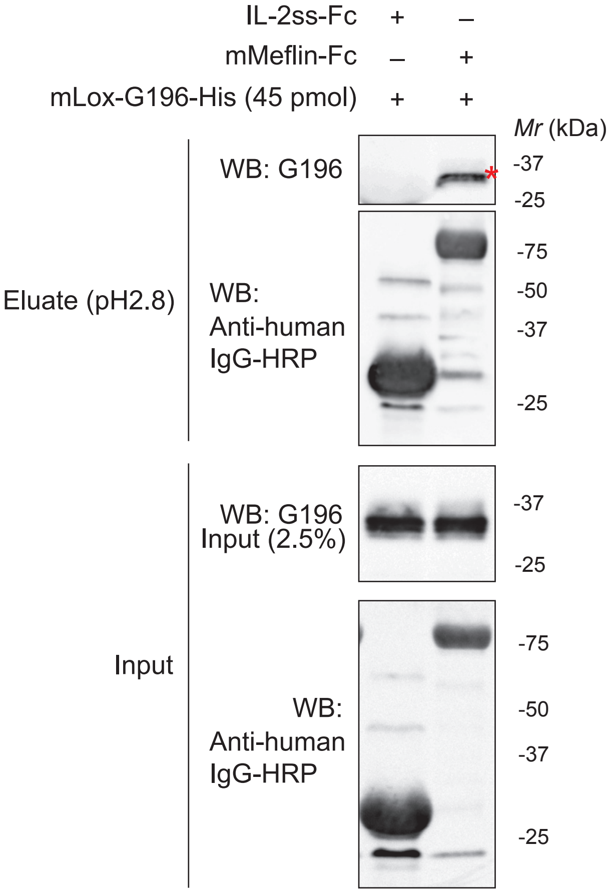


**f**

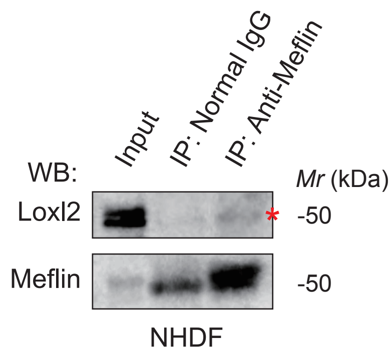


**Supplementary Fig 6**

**Supplementary Fig 6.** Identification of Lox as a Meflin-interacting protein.

**(a, b)** Identification of Meflin-interacting proteins in mouse MSCs. Mouse MSCs were infected with retroviruses encoding either signal sequence (SS)-Flag-HA-mMeflin or GFP, followed by IP with anti-HA antibodies. The Meflin immunoprecipitate was then eluted with HA peptide and analyzed by silver staining **(b)** and mass spectrometry.

**(c)** List of proteins detected in the Meflin immunoprecipitate. The list includes only secreted or membrane proteins detected in the Meflin immunoprecipitate given that Meflin is a GPI-anchored membrane or secreted protein. ISLR, which is the official symbol for Meflin, was identified in the Meflin immunoprecipitate as expected (asterisk). Lox (UniProt Knowledgebase entry name LYOX_MOUSE) was identified as the highest score protein in the Meflin immunocomplex (arrow).

**(d)** Meflin interacts with Loxl2. The indicated plasmids were cotransfected into 293FT cells, followed by IP with anti-HA antibodies and WB analysis. The data showed that Meflin interacted with human Loxl2 (asterisk).

**(e)** Direct interaction of Meflin and Lox proteins. Purified mMeflin (1–399 aa)-Fc (constant region of IgG1) and Fc fused with the signal sequence of IL-2 (IL-2ss) were mixed with recombinant mouse Lox tagged with the G196 and His epitopes (mLox-G196-His, 45 pmol). Samples were then washed, eluted, and subjected to WB analysis with the indicated antibodies. An asterisk indicates Lox proteins that bound to Meflin-Fc.

**(f)** Interaction of endogenous Meflin and Loxl2 proteins. Cell lysates prepared from the primary cultured human fibroblasts (NHDFs) were immunoprecipitated with anti-Meflin antibodies, and eluates were analyzed by WB using the indicated antibodies. An asterisk indicated Loxl2 co-precipitated with Meflin.

**a**

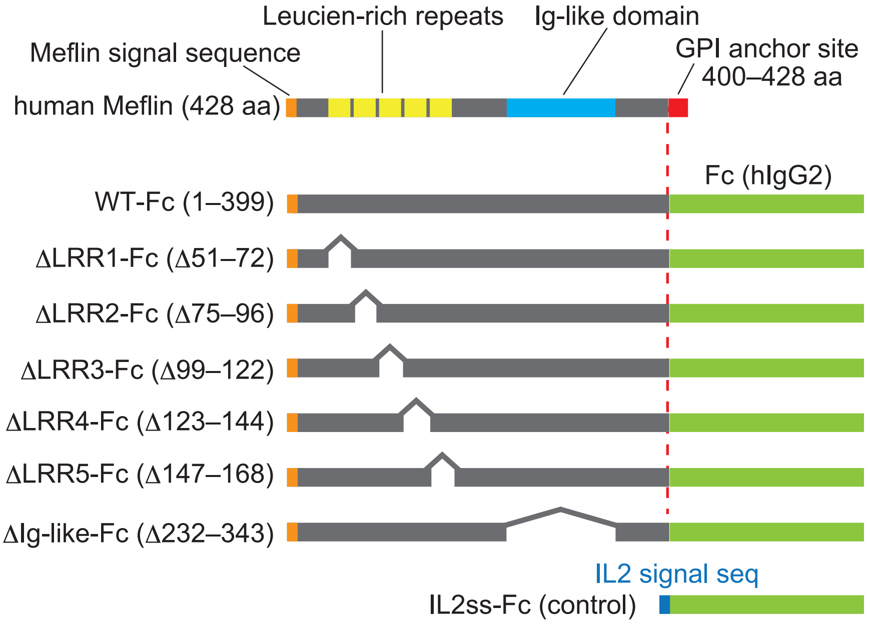


**b**

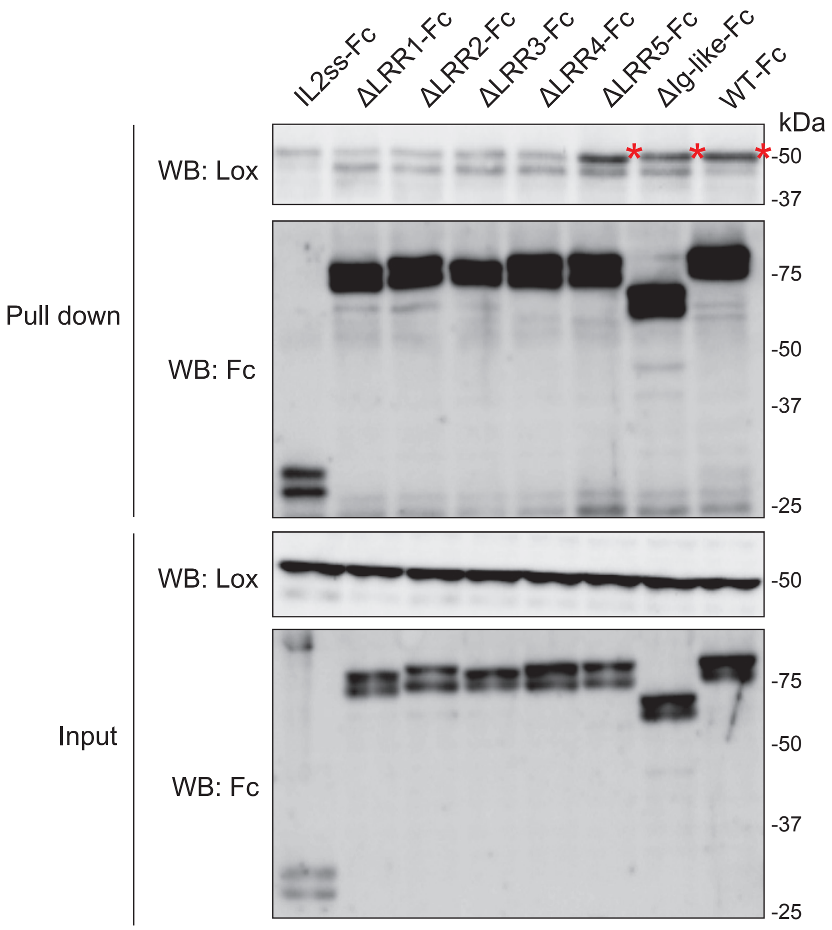


**Supplementary Fig 7**

**Supplementary Fig 7.** Requirement of the leucine-rich repeat (LRR) domain of Meflin for interaction with Lox.

**(a)** Schematic illustration of full-length human Meflin and its mutants lacking one LRR of the LRR domain, consisting of LRR1–5, and the immunoglobulin (Ig)-like domain. cDNAs encoding all mutants fused with IgG1 Fc were subcloned into the pRP expression vector. Fc fused with IL-2ss was used as a control.

**(b)** Weak interaction of Meflin mutants lacking LRR1, 2, 3, and 4 domains with Lox. The indicated plasmids were cotransfected into 293FT cells, followed by precipitation with protein A agarose and elution with a sodium dodecyl sulfate sample buffer. The eluted proteins were analyzed using the indicated antibodies. Note that ΔLRR1-Fc, ΔLRR2-Fc, ΔLRR3-Fc, and ΔLRR4-Fc mutants exhibited very weak interactions with Lox, in contrast to ΔLRR5-Fc and ΔLRR4-Fc mutants and WT-Fc (asterisks).

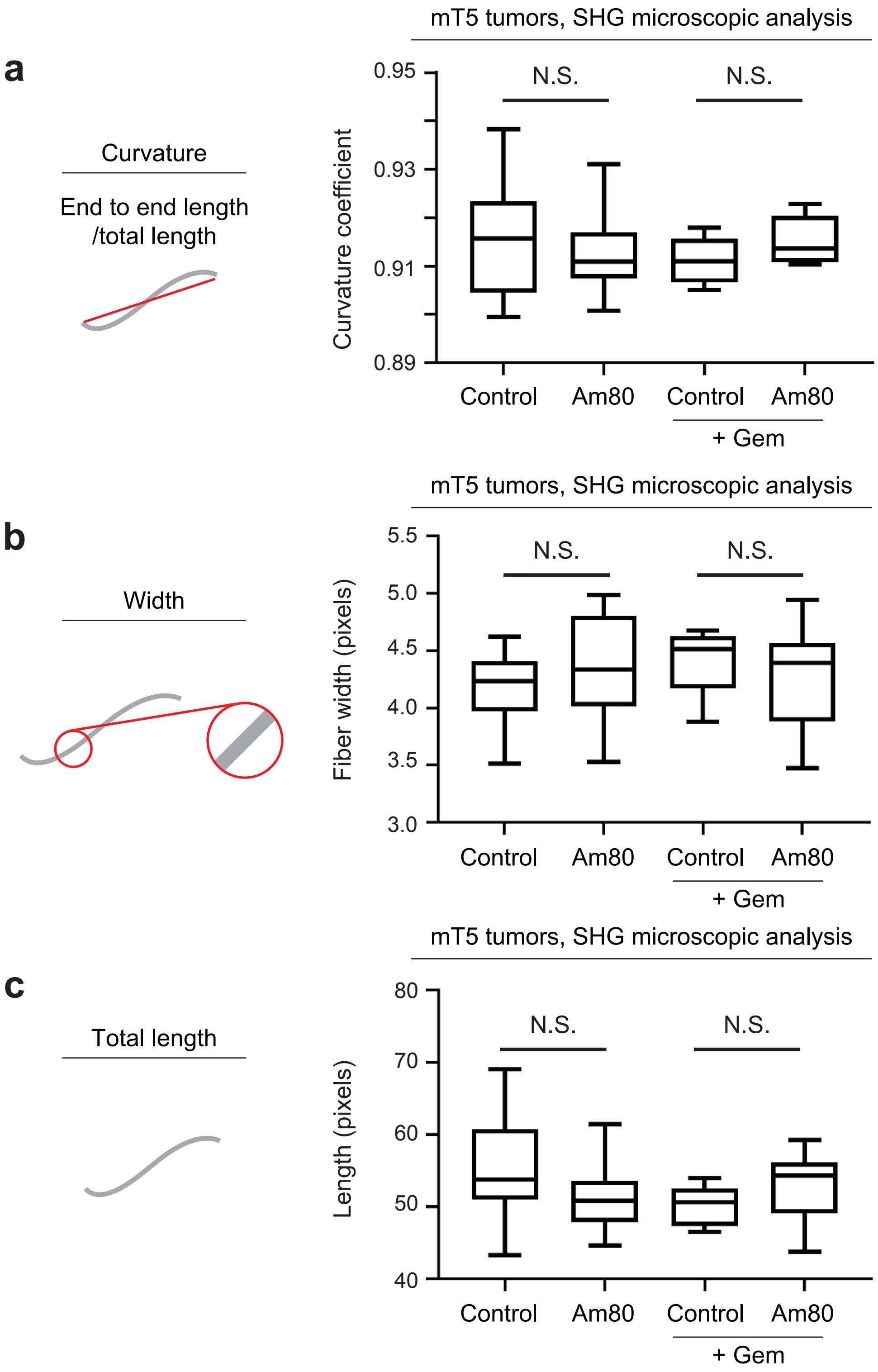


**Supplementary Fig 8**

**Supplementary Fig 8.** Measurement of collagen alignment in the stroma of mT5 tumors by SHG microscopy.

**(a–c)** Eight to sixteen randomly selected images from tissue sections of control tumors and those treated with Am80 in the presence or absence of gemcitabine treatment were analyzed for collagen curvature **(a)**, width **(b)**, and total length **(c)** by SHG microscopy.

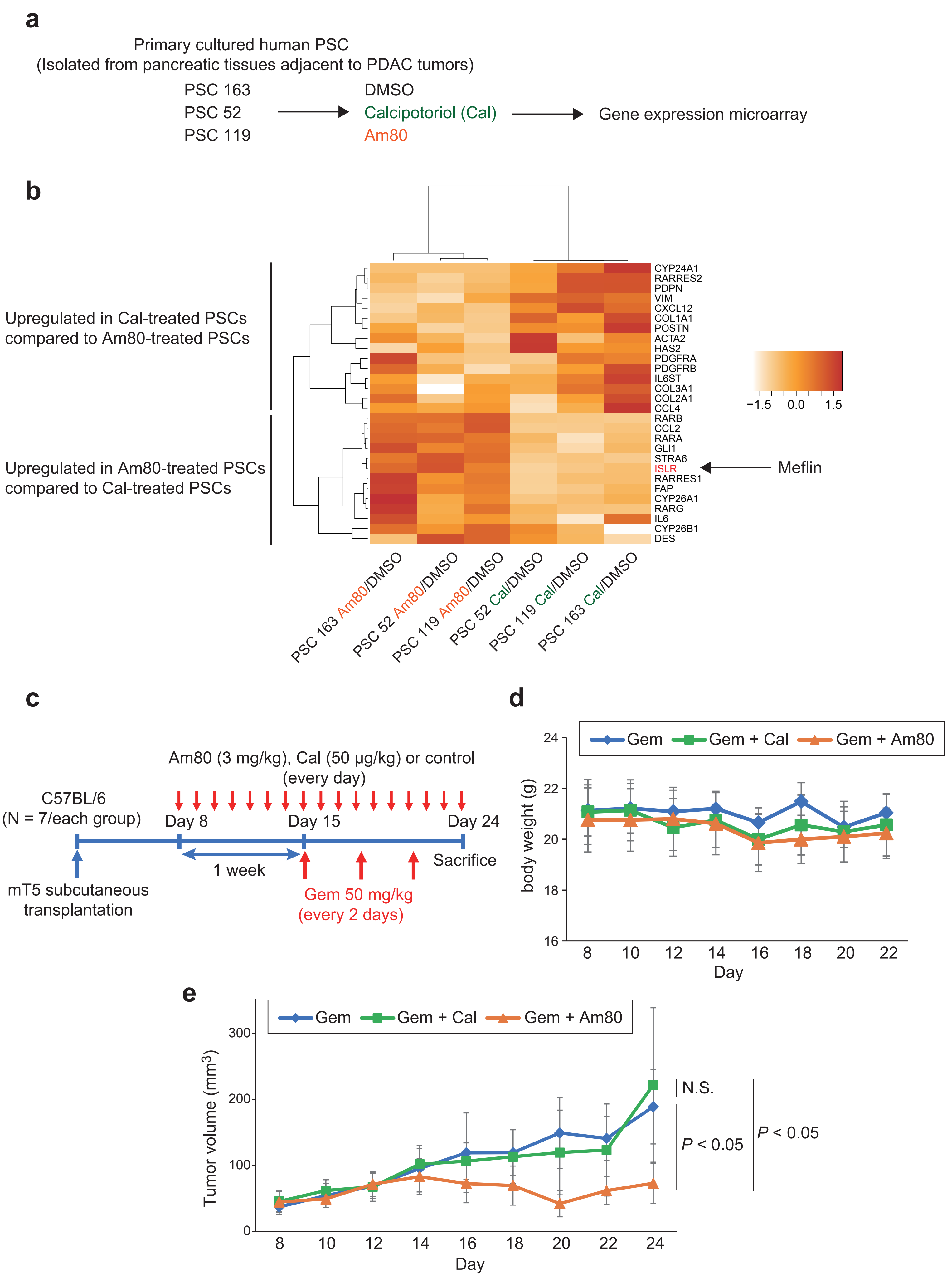


**Supplementary Fig 9**

**Supplementary Fig 9.** Differential effects of Am80 and calcipotriol administration on gene expression in CAFs and tumor progression in a PDAC xenograft model

**(a)** Primary cultured human PSCs isolated from three patients with PDAC (163, 52, and 119) were administered with DMSO (1 μM), calcipotriol (Cal, 1 μM), and Am80 (1 μM) for 48 h. Total RNA was then extracted, and gene expression microarray analysis was performed.

**(b)** Expression values of CAF marker genes and genes involved in vitamin D or A signaling in Cal- and Am80-treated PSCs were divided by the corresponding gene values in DMSO-treated PSCs, and comparisons between the two groups (Am80/DMSO versus Cal/DMSO) were performed. Genes that were reproducibly upregulated or downregulated in at least two PSC lines compared with the other group were extracted and shown as a heatmap. Data are represented as $\log_2$ fold changes between Cal/DMSO and Am80/DMSO groups.

**(c)** WT female mice (P42) were subcutaneously implanted with mT5 cells ($1 \times 10^6$ cells/mouse) and maintained until the tumors reached 50–100 $mm^3$ in volume. Oral administration of calcipotriol (50 μg/kg, 0.5% CMC solution), Am80 (3 mg/kg, 0.5% CMC solution), or 0.5% CMC solution (control) was performed for 16 consecutive days (days 8–24). Mice were administered Gem from day 15 three times every 3 days.

**(d, e)** Measurement of body weights **(d)** and tumor volumes **(e)** for the indicated groups during the observation period.

### Supplementary Methods

#### ISH

All ISH analyses were performed on FFPE human and mouse tissue samples using RNAscope technology (RNAscope 2.5 HD Detection Kit; Advanced Cell Diagnostics) and custom-designed probes following the manufacturer's instructions. Briefly, tissue sections were baked on a dry oven (HybEZ II Hybridization System; Advanced Cell Diagnostics) at 60°C for 1 h, deparaffinized, and incubated with $H_2O_2$ solution (Pretreat 1 buffer) for 10 min at room temperature. Slides were boiled in a target retrieval solution (Pretreat 2 buffer) for 15 min and incubated with a protease solution (Pretreat 3 buffer) for 30 min at 40°C. Slides were then incubated with the relevant probes for 2 h at 40°C and successively incubated with Amp1–6 reagents (Advanced Cell Diagnostics). Staining was visualized with 3,3-diaminobenzidine, followed by counterstaining with hematoxylin. The RNAscope probes used in this study were human Meflin (*ISLR*; NM_ 005545.3, region 275–1322, cat. no. 455481), mouse Meflin (*Islr*; NM_012043.4, region 763–1690, cat. no. 450041), and mouse *Acta2* (NM_007392.3, region 41–1749, cat. no. 319531).

#### Xenograft tumor models

In the transplantation models of mouse mT5 PDAC cells and human BxPC3 cells, $1 \times 10^6$ cells were transplanted subcutaneously into the backs of C57BL/6 adult WT, Meflin-KO, or nude mice. For the experiment shown in **Fig. 3**, Am80 (20 mg/mL) in 0.5% carboxymethylcellulose (CMC) solution (Wako Chemicals, Osaka, Japan) was orally administered daily at doses of 0, 0.1, 0.5, 1, or 3 mg/kg. For experiments in **Figs. 4** and **5**,

C56BL/6 adult WT or Meflin-KO mice were randomized into the following treatment groups: saline, Gem (50 mg/kg, i.p.), and Am80 (3 mg/kg/day, oral administration) plus Gem (50 mg/kg, i.p.). For comparison of the effects of Am80 and calcipotriol on the efficacy of Gem in **Supplementary Fig. 9**, we began Am80 and calcipotriol treatment on day 8, followed by Gem treatment on day 15.

In all experiments, mice were euthanized around day 20 after tumor implantation, after which tumors were harvested, sliced, and flash-frozen in liquid nitrogen or immediately fixed in a formalin solution for subsequent histological analysis. Tumor sizes were measured on the indicated days with calipers, and tumor volumes were calculated using the formula $X^2 \times Y \times 0.5$, where X is the smaller diameter and Y is the larger diameter.

In the experiment shown in **Supplementary Fig. 5**, KPC mice with PDAC were enrolled in the study based on tumor size, as described previously (1). Enrollment was restricted to mice with tumors greater than 10 mm in diameter, as determined by high-resolution ultrasound imaging (Vevo 2100 ultrasound system; Fujifilm Visual Sonics). Suitable mice were assigned to the following treatment groups: Gem or Gem plus Am80. KPC mice were orally administered saline or Am80 (20 mg/mL, 0.5% CMC solution) at 3 mg/kg from day 1, followed by i.p. injection of Gem (10 mg/mL, saline) at a dose of 50 mg/kg three times every 3 days. The longest diameter of the largest tumor between the observed tumors was evaluated.

### Immunohistochemistry

FFPE tissue sections were deparaffinized, followed by antigen retrieval by boiling the samples in Target-Retrieval Solution (Dako) at pH 6, 7, or 9 for 30 min and conventional staining procedures, as described previously (20).

**Antibodies**

The antibodies used in this study included rabbit polyclonal anti-Meflin antibodies (Atlas antibody; cat. no. HPA050811, 1:1000), mouse monoclonal anti-α-SMA antibodies (Dako; clone 1A4, 1:1000), rabbit polyclonal anti-Ki-67 antibodies (Novus Biologicals; cat. no. NB600-1252, 1:100), polyclonal anti-GFP antibodies (MBL; cat. no. 598, 1:1000), mouse monoclonal β-actin antibodies (Sigma, St. Louis, MO, USA; cat. no. A5316, 1:2000), rat monoclonal anti-CD31 antibodies (Dianova; cat. no. DIA-303, 1:50), rabbit polyclonal anti-RFP antibodies (Rockland; cat. no. 600-401-379), rabbit monoclonal anti-Lox antibodies (Abcam, Cambridge, UK; cat. nos. ab174316 and ab174316, 1:1000), mouse monoclonal anti-HA antibodies (Sigma; cat. no. A2095, 1:30), and rabbit polyclonal anti-Loxl2 antibodies (GeneTex; cat. no. GTX105085, 1:1000). For immunoprecipitation of endogenous Meflin, we used rat monoclonal anti-Meflin antibodies generated in our laboratory.

**Culture of cell lines and isolation of human PSCs**

The mouse PDAC cell line mT5 was generously provided by David Tuveson (Cold Spring Harbor Laboratory) and Chang-Il Hwang (UC Davis College of Biological Sciences, Davis, CA). BxPC3 and C3H10T1/2 cell lines were purchased from the American Type Culture

Collection (Manassas, VA, USA). C57BL/6J bone marrow-derived MSCs were purchased from Cyagen Biosciences. fHDF/TERT166 cells were purchased from Evercyte. NHDFs were purchased from Lonza. All cell lines were authenticated by routine morphological and growth analyses and routinely screened for *Mycoplasma* contamination by 4',6-diamidino-2-phenylindole staining.

Human PSCs were isolated from pancreatic tissues adjacent to primary PDAC tumors during surgery at Tohoku University Hospital, as described previously (2, 3). Immortalized human PSCs were established by retrovirus-mediated introduction of the simian virus 40 (SV40) large T antigen (3). This study was approved by the Ethics Committees of Tohoku University (approval number 2017-1-280) and Nagoya University Graduate School of Medicine (approval number 2015-0254). All cells were cultured in Dulbecco's modified Eagle's medium (DMEM; Nacalai Tesque, Japan) supplemented with 10% fetal bovine serum (FBS).

**Generation of SeV18+mMeflin/TSΔF**

An insert sequence containing the open reading frames of the mouse Meflin (mMeflin, *Islr*) gene or DasherGFP (DGFP) gene was constructed by PCR from cDNAs using NotI-tagged gene-specific forward and reverse primers containing SeV-specific transcriptional regulatory signal sequences. Amplified fragments were inserted into the 18+ region of the plasmid containing the SeV/TSΔF vector sequence, producing pSeV18+mMeflin-DGFP/TSΔF or pSeV18+DGFP/TSΔF. Recovery and propagation of SeV18+mMeflin-DGFP/TSΔF or SeV18+DGFP/TSΔF were performed as follows. First, 293T cells were transfected with pSeV18+mMeflin-DGFP/TSΔF or pSeV18+DGFP/TSΔF and pCAGGS plasmids carrying the T7 RNA polymerase and NP, P, F5R, and L genes. Cells were maintained in DMEM supplemented with 10% heat-inactivated FBS and cultured for 1–3 days to generate the seed of the SeV18+mMeflin-DGFP/TSΔF vector or SeV18+DGFP/TSΔF vector. The seeds were cloned and propagated using SeV F-expressing LLC-MK2/F7/A cells (4) in MEM containing trypsin (2.5 μg/mL). The titer (cell infectious units/mL) of the recovered SeV18+mMeflin-DGFP/TSΔF vector or SeV18+DGFP/TSΔF vector was determined by the immunostaining method using anti-SeV rabbit polyclonal sera as described previously (5).

To test the effects of Meflin transduction on the expression of CAF marker genes, mouse MSCs and C3H10T1/2 cells were transduced with the SeV vectors at 20 MOI, followed by culture in DMEM/10% FBS at 35°C in 5% $CO_2$ for 16 h. The medium was replaced with DMEM/10% FBS after 24 h of infection, and the cells were cultured at 37°C in 5% $CO_2$ thereafter.

**Western blot analysis**

Cells were lysed in lysis buffer containing 30 mM Tris-HCl (pH 7.4), 120 mM NaCl, 1 mM ethylenediaminetetraacetic acid (EDTA), 1% Triton-X 100, 20 mM β-glycerophosphate, and 1 mM p-(amidinophenyl) methanesulfonyl fluoride hydrochloride supplemented with Complete Protease Inhibitor (Roche) and PhosSTOP Phosphatase Inhibitor cocktails (Roche). Lysates were clarified by centrifugation at 12,000 × *g* for 10 min at 4°C, followed by the addition of sodium dodecyl sulfate (SDS) sample buffer (10 mM Tris-HCl, 2% SDS, 2 mM EDTA, 0.02% bromophenol blue, 6% glycerol; pH 6.8) and separation by SDS-polyacrylamide gel electrophoresis (PAGE). Proteins were transferred to nitrocellulose membranes, blocked in 5% milk in phosphate-buffered saline (PBS) containing 0.05% Tween 20, incubated with primary antibodies, and detected by horseradish peroxidase (HRP)-conjugated secondary antibodies (Dako).

**Quantitative polymerase chain reaction (qPCR)**

For analysis of gene expression in primary cultured mouse MSCs, human PSCs, and C3H10T1/2 cells, total RNA was extracted using RLT buffer (Qiagen, Valencia, CA, USA) and the RNeasy Mini Kit (Qiagen) and was treated with RNase-free DNase (Qiagen) according to the manufacturer's instructions. Purified RNA samples were reverse-transcribed using ReverTra Ace (Toyobo, Tokyo, Japan) with oligo dT and random primers. qPCR of the generated cDNAs was performed with TaqMan Gene Expression Master Mix (Applied Biosystems, Foster City, CA, USA) on an Mx3005P thermal cycler (Agilent Technologies). TaqMan probes and primers for mouse Meflin (*Islr*; Mm01700423_m1), mouse α-SMA (*Acta2*; Mm00725412_s1), mouse IL6 (*Il6*; Mm00446190_m1), mouse Col1a1 (*Col1a1*; Mm00801666_g1), mouse Col3a1 (*Col3a1*; Mm01254476_m1), and mouse β-actin (*Actb*; Mm04394036_g1) were purchased from Life Technologies and used according to the manufacturer's instructions. Cycling conditions were as follows: 95°C for 10 min, 40 cycles of 95°C for 15 s, and then 60°C for 1 min, followed by one cycle of 95°C for 10 s. The data were analyzed using the $2^{-\Delta\Delta ct}$ method and normalized to the *Gapdh* control.

**Screening of a nuclear receptor ligand library**

Primary cultured human PSCs isolated from pancreatic tissue adjacent to PDAC or mouse MSCs (Cyagen) were plated in 6-well plates and allowed to reach superconfluence. The compounds included in the nuclear receptor ligand library (74 compounds; ENZO Life Sciences; cat. no. BML-2802-0100) were added to duplicate wells of the 6-well plates at a final concentration of 1 μM in 0.1% dimethyl sulfoxide (DMSO). The cells were cultured for 48 h, followed by extraction of total RNA using an RNeasy Plus Mini Kit (Qiagen) and analysis of Meflin (*Islr*) expression by qPCR.

**Reagents for animal studies**

Concentrated stock solutions (10 mM) of Am80 (cat. no. 3507; Tocris) and calcipotriol (cat. no. 2700; Tocris) were prepared in DMSO and stocked at −20°C, protected from light, for up to 1 month. The stock solutions were diluted to the appropriate dose just before use. Gemcitabine (Gemzar; Eli Lilly) was dissolved in 0.9% sterile saline just before each experiment and administered in a volume of 0.1 mL/mouse by i.p. injection.

**Sample preparation and liquid chromatography tandem mass spectrometry (LC-MS/MS) analysis**

LC-MS/MS was performed as described by Bapiro *et al.* (6). Weighed tumor samples (10 mg) were disrupted using a Multi-bead Shocker (Yasui Kikai, Osaka, Japan) according to the manufacturer's instructions and mixed with 200 μL ice-cold 50% v/v acetonitrile (cat. no. 01031-1B; Kanto Chemical Co., Inc.) containing 25 μg/mL tetrahydrouridine (CAS18771-50-1; Merck). After short-term storage at −80°C, the samples were thawed on ice and homogenized with a single 3-s pulse in a Vibra-CellTM Ultrasonic Liquid Processors (cat. no. VC501; Sonics & Materials, Inc.) set to amplitude 30. An aliquot (50 µL) of the homogenate was added to a microfuge tube with 200 µL ice-cold acetonitrile (50% v/v) containing 250 ng/mL 2′-deoxy-2′,2′-difluorocytidine-13C, 15N2 dFdC (cat. no. G305002; TRC) as an internal standard. Vortex mixing was followed by centrifugation at 20,000 × *g* for 25 min, and the resulting supernatant was filtered using a 0.1-µm filter and evaporated to dryness in a Speedvac. The residue was reconstituted in 10 mM ammonium acetate (pH 10.0) and desalted using GL-Tip GC (GL Science). Desalted samples were subjected to LC-MS (Dionex Ultimate 3000 HPLC system equipped with an autosampler and an EXACTIVE Plus mass spectrometer [Thermo] fitted with a Hypercarb column [5 µm, 2.1 × 100 mm; Thermo]) using (A) 10 mM aqueous ammonium acetate (pH 10.0) and (B) acetonitrile as mobile phase. The autosampler and column temperatures were maintained at 10 and 30°C, respectively. The gradient program was as follows: flow rate of 300 µL/min; 95% A for 2 min, decrease to 80% at 0.2 min and held for 5.6 min, increase to 95% over 0.2 min, held at 95% for 7 min (total run-time of 15 min). Samples were detected using electrospray ionization in positive ion mode. dFdC was quantified against the corresponding isotope-labeled dFdC as an internal standard. Data were acquired and analyzed with Xcalibur ver. 2.2 software (Thermo).

**Plasmids**

Cloning of mouse Meflin (mMeflin) cDNA was described previously (20, 21). cDNAs for mouse Lox (mLox) and human Loxl2 (hLoxl2) were obtained from Origene (Lox-Myc-DDK-mLox; cat. no. MR206463) and Open Biosystems (clone ID: 3347512), respectively, and subcloned into the pEF5/FRT/V5 D-TOPO vector (Thermo Fisher Scientific). To generate mMeflin fused with the constant region (Fc) of human Ig, mMeflin (amino acid residues 1–399), which lacks a carboxyl-terminal GPI anchor site, was amplified by PCR and inserted into the pFUSE-hIgG2-Fc1 vector (InvivoGen). mMeflin cDNA was also inserted into the retrovirus expression vector pRetroQ (Clontech) with the SS-Flag-HA tag at the amino-terminus (pRetroQ-SS-HA-mMeflin). Generation of human Meflin and its deletion mutants fused with the Fc of human Ig was commercially produced by Vector Builder (Guangzhou, China).

#### Identification of Meflin-interacting proteins by MS

Primary cultured mouse MSCs infected with retroviruses encoding either GFP or SS-FH-mMeflin were lysed in a buffer (50 mM Tris-HCl, 150 mM NaCl, 2 mM EDTA, 0.1% NP-40, pH 7.5), immunoprecipitated with monoclonal anti-HA-agarose (Sigma; cat. no. A2095, clone HA-7), and eluted with HA peptide (Sigma; cat. no. I2149, 1 mg/mL). Whole immunoprecipitates were then digested with Trypsin Gold (Promega, Madison, WI, USA) for 16 h at 37°C and analyzed on a Q Exactive mass spectrometer (Thermo Fisher Scientific).

#### Immunoprecipitation (IP)

For IP, cells were lysed in buffer containing 20 mM Tris-HCl (pH 7.4), 120 mM NaCl, 1 mM EDTA, and 1% Triton X-100 supplemented with cOmplete Protease Inhibitor cocktail (Roche) and PhosSTOP phosphatase inhibitor cocktail (Roche). Lysates were cleared by centrifugation at 12,000 × *g* for 10 min at 4°C, and IP was carried out using the indicated antibodies and protein A/G beads (Sigma).

#### *In vitro* binding assay

Expi293F suspension cells (Thermo Fisher Scientific) cultured in 30 mL medium were transduced with mMeflin (1–399 aa) fused with the Fc region of human IgG2 (mMeflin-Fc) or control Fc with the SS of IL-2 (IL-2ss-Fc) inserted into an expression vector pEB Multi-Hyg (Wako Pure Chemicals), followed by culture for 5 days. mMeflin-Fc and IL2ss-Fc proteins expressed in the medium were purified by batch affinity purification using Ab-capcher ExTra (ProteNova, Japan), followed by extensive washing with PBS. The purified mMeflin-Fc and IL-2ss-Fc were mixed with 45 pmol recombinant mLox (mLox-G196-His) purified as described above in a binding buffer (ProteNova) for 3 h at 4°C, followed by washing with PBS and elution with an elution buffer (pH 2.8; ProteNova). The eluted samples were neutralized with neutralization buffer (ProteNova) and subjected to western blot analysis.

#### Pull-down assays

HEK293T cells were transfected with mLox and the indicated Meflin mutants fused with Fc at the C-terminus and cultured for 48 h. Cells were then lysed with lysis buffer and clarified by centrifugation at 12,000 × *g* for 10 min at 4°C. The soluble supernatant was incubated with protein A beads (Sigma) for 8 h at 4°C. RNase (10 µg/mL; Sigma) was added to lysis buffer to reduce nonspecific binding of proteins to the beads. The beads were washed extensively with lysis buffer before elution of the purified Fc fusion proteins and their binding proteins with SDS sample dilution buffer. Aliquots of the eluates were subjected to SDS-PAGE, followed by western blot analysis using the indicated antibodies.

**Generation of cells stably expressing mMeflin**

For generation of an isogenic cell line stably expressing mMeflin, we utilized the Flp-In system (Thermo Fisher Scientific). Briefly, pOG44 (Thermo Fisher Scientific) and pcDNA5/FRT (Thermo Fisher Scientific) harboring mMeflin were cotransfected into Flp-In-293 cells (Thermo Fisher Scientific). Twenty-four hours after transfection, the cells were trypsinized and transferred into 5 × 10-cm dishes, followed by selection on hygromycin (180 µg/mL) for 7–10 days. Pools of all of foci, which represented isogenic populations, were used for subsequent experiments.

**Lox activity assay**

Culture supernatants of control Flp-In-293 cells or those stably expressing mMeflin were collected and centrifuged at 13,000 × *g* for 5 min at 4°C. The samples were then equilibrated to room temperature and mixed with recombinant human Loxl2 (R&D Systems; cat. no. 2639-AO) at the indicated concentrations, followed by addition of a proprietary Lox substrate that releases hydrogen peroxide upon transformation by Lox activity or its equivalent present in the samples (Lysyl Oxidase Activity Assay Kit; Abcam). Hydrogen peroxide was detected using a proprietary red fluorescence substrate for

HRP-coupled reactions. The fluorescence was measured at Ex/Em = 540/590 nm in a fluorescence microplate reader (Powerscan4; BioTek).

**SHG imaging and analysis**

All samples were imaged on a custom-built, multiphoton microscope, consisting of a Nikon TE300 inverted microscope, Coherent Chameleon XR laser path, and Hamamatsu H7422P-40 photomultiplier with a 40× objective. SHG was collected using laser excitation at 890 nm with emission filtration using a Semrock Brightline 445/20nm filter. For each sample, 5 image plane z-stacks were collected using a 2-µm step size and projected using maximum intensity. Collagen fibers were identified from SHG images using ct-FIRE (loci.wisc.edu/software/ctFIRE, v.2.0b). Further analysis of fiber to fiber thickness, straightness, and orientation was completed using CurveAlign (loci.wisc.edu/software/curvealign,v.4.0b).

**Measurement of Young's modulus of mT5 xenograft tumors**

mT5 subcutaneous xenograft tumors developed in the back skin of mice 15 days after transplantation were harvested and separated from the panniculus carnosus muscle. The tumors were embedded in an autopolymerizing resin (Quick Resin, Shofu Inc., Japan), followed by flipping upside down such that the flat surfaces of the tumors that had contact with the fascia of the panniculus carnosus muscle were facing upward. The Young's modulus of the tumors was measured using Softgram (Shinko Denshi Co., Ltd., Japan), which is a device for the determination of the stiffness of biomaterials based on the Hertz elastic contact theory.

### cDNA gene expression microarray analysis

Primary cultured PSCs were plated in 6-well plates and allowed to become superconfluent. Next, AM80, calcipotriol, or DMSO was added to PSCs at a final concentration of 1 μM in 0.01% DMSO, and cells were cultured for 48 h, followed by extraction of total RNA with an RNeasy Plus Mini Kit (Qiagen). Cyanine-3 (Cy3)-labeled cRNA was prepared from 500 ng RNA using a Quick Amp Labeling Kit (Agilent) according to the manufacturer's instructions. Fragmentation of Cy3-labeled cRNA and hybridization to an Agilent SurePrint G3 Mouse GE 8x60K Microarray were performed by Takara Bio (Yokkaichi, Japan).

### Statistical analyses

Data are presented as means ± standard deviations of the means. Means of 2 groups were compared using Mann-Whitney U tests or Brunner-Munzel tests. For multiple comparisons, we used analysis of variance with subsequent Bonferroni correction. Statistical analyses were conducted using GraphPad Prism 7.00 (GraphPad Software Inc.).
